# Selective vulnerability of supragranular layer neurons in schizophrenia

**DOI:** 10.1101/2020.11.17.386458

**Authors:** Mykhailo Y. Batiuk, Teadora Tyler, Shenglin Mei, Rasmus Rydbirk, Viktor Petukhov, Dora Sedmak, Erzsebet Frank, Virginia Feher, Nikola Habek, Qiwen Hu, Anna Igolkina, Lilla Roszik, Ulrich Pfisterer, Zdravko Petanjek, Istvan Adorjan, Peter V. Kharchenko, Konstantin Khodosevich

**Author notes:** correspondence should be addressed to: IA PVK and KK. authors have contributed equally (Mykhailo Y. Batiuk, Teadora Tyler).

## Abstract

Schizophrenia is one of the most wide-spread mental brain disorders with complex and largely unknown etiology. To characterize the impact of schizophrenia at a cellular level, we performed single nucleus RNA sequencing of >190,000 neurons from the dorsolateral prefrontal cortex of patients with schizophrenia and matched controls (7 vs 11, respectively). In addition, to correlate data with cortical anatomy, >100,000 neurons were analyzed topographically by immunohistochemistry in an extended cohort of cases with schizophrenia and controls (10 vs 10). Compositional analysis of RNA sequencing data revealed reduction in relative abundance across all families of GABAergic neurons and a concomitant increase in principal neurons, which was most pronounced for supragranular subtypes (layers 2-3). Moreover, supragranular subtypes of GABAergic interneurons showed most dramatic transcriptomic changes. These results were substantiated by histological analysis, which revealed a reduction in the density of calretinin, calbindin and parvalbumin GABAergic interneurons particularly in layer 2. Common effect of schizophrenia on supragranular neuronal networks was underlined by downregulation of protein processing genes and upregulation of neuronal development/plasticity genes across supragranular subtypes of principal neurons and GABAergic interneurons. *In situ* hybridization and spatial transcriptomics further confirmed supragranular layer neuron vulnerability, revealing complexity of schizophrenia-affected cortical circuits. These point towards general network impairment within supragranular layers being a core substrate associated with schizophrenia symptomatology.

---

Schizophrenia is a severe mental brain disorder, which affects 20-60 million people worldwide. Due to high complexity of neuronal circuits that underlie cognitive abnormalities in schizophrenia, the etiology of schizophrenia is poorly understood^1,2^. The previous work has been mainly focused on analysis of selected types of neurons, e.g. expressing parvalbumin (PV), showing decrease in their density and change in gene expression^3–6^. Some effort has been made for less biased characterization of schizophrenia-associated changes in neurons using bulk transcriptomics in the human brain. While these studies identified important generalized effects of schizophrenia on cortical tissue^3,7–10^, they lacked the resolution to answer questions what neurons might be most affected in schizophrenia and how schizophrenia-associated gene expression could contribute to abnormal neuronal circuitry in schizophrenia. Thus, to reveal impact of schizophrenia on whole diversity of neuronal subtypes in the cortex, we used single-nucleus RNA sequencing (snRNA-seq) for a cohort of schizophrenia patients and controls (7 and 11, respectively) and examined the detailed transcriptional states of neurons in the dorsolateral prefrontal cortex (DLPFC) from Brodmann area 9 (BA9). The BA9 was chosen based on the functional and anatomical data from schizophrenia patients showing abnormality in brain activity, neuronal morphology and gene expression in this area relative to matched control brains^11–13^. Importantly, our sample cohort was formed by strictly limiting the potential confounders, including the age of sample donors (the vast majority of samples were from 55-65 years old subjects), their post-mortem intervals (PMI, below 20 hours) and other confounders (Supplementary Table 1). Furthermore, the schizophrenia and control groups were balanced with respect to multiple other characteristics, including sex, medication, smoking etc (Extended Data Fig. 1, and Methods).

For each sample, we microdissected cortical columns containing all layers, sorted for neuronal nuclei based on the expression of the neuronal marker NeuN (Fig. 1a, Extended Data Fig. 2a) and performed snRNA-seq using 10X Chromium assay. In total, we sequenced 191,407 nuclei, of which 174,560 (67,907 from schizophrenia and 106,653 control nuclei) passed quality control analysis and doublet filtering (Extended Data Fig. 2b-e, Supplementary Table 1).

**Figure 1.**
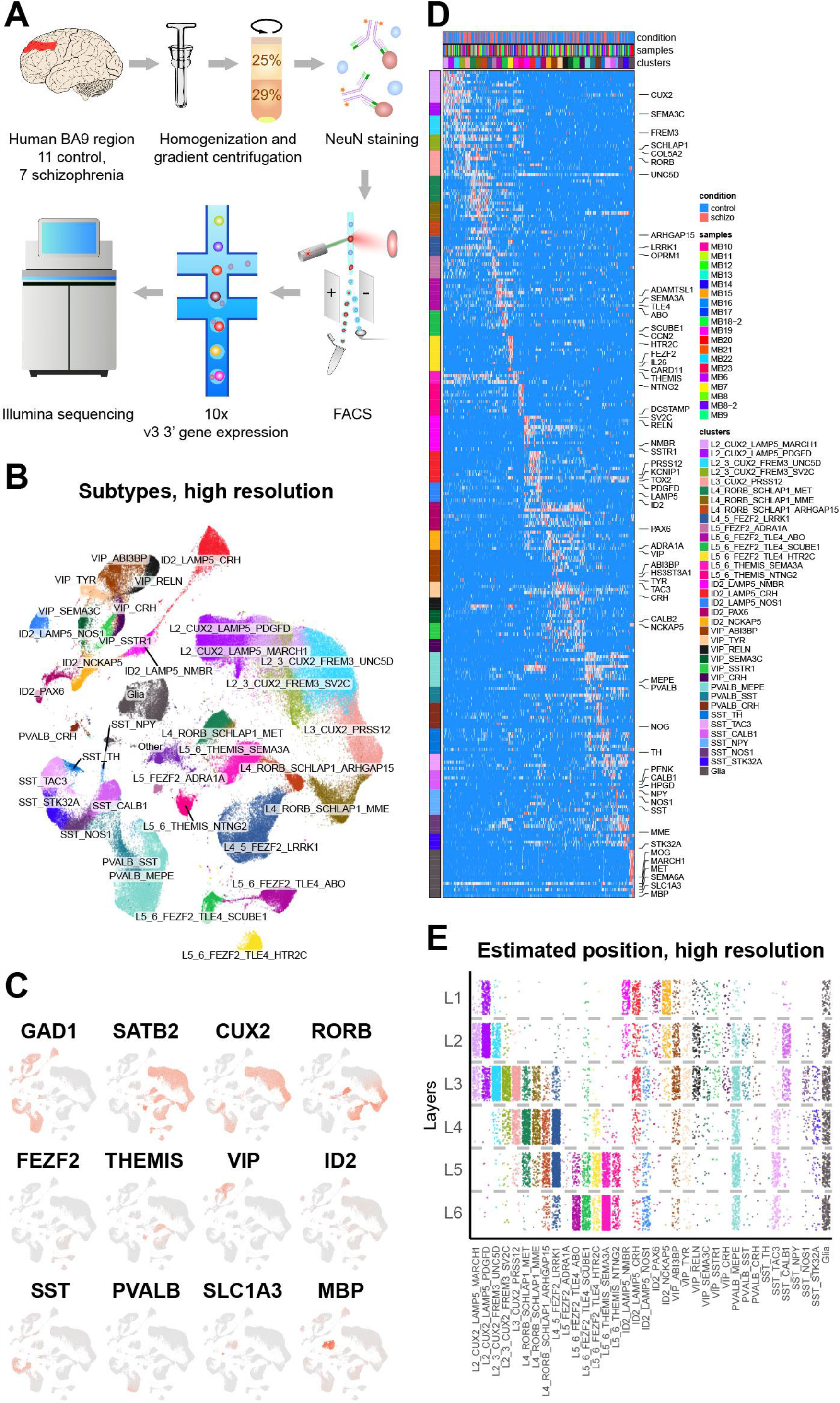
snRNA-seq analysis of the DLPFC from patients with schizophrenia and matched controls. A: Experimental design of the snRNA-seq measurements. B: UMAP representation of the measured nuclei, colored by subtypes. C: Expression of marker genes for major cortical cell types, visualized on the UMAP embedding: GABAergic interneurons, principal neurons and glia; additional markers distinguish families of GABAergic interneurons and principal neurons. D: Heatmap showing expression of markers for specific neuronal subtypes. E: Estimated cortical layer positions of the neuronal subtypes.

To annotate the neuronal populations in a consistent manner, different samples were first aligned in a way that minimized the impact of the inter-individual variation, and clusters of nuclei were determined uniformly across the entire sample collection^14^. The clusters were annotated based on known layer-specific genes for principal neurons, subtype-specific genes for GABAergic interneurons from previous studies^15,16^, as well as cluster-specific markers (Supplementary Table 2). The subpopulations were annotated in a hierarchical manner, with the medium resolution denoting neuronal subfamilies, and high resolution representing specific subtypes (Supplementary Table 2). At the highest resolution we distinguished a total of 15 principal neuron and 20 GABAergic interneuron transcriptomic subtypes (Fig. 1a-d, Extended Data Fig. 3a,b, Supplementary Table 2). There was an even representation of each subtype from multiple control and schizophrenia samples, and every subtype was present in at least 75% of the samples (Extended Data Fig. 3c-f). Age, PMI, sex and other sample characteristics were evenly distributed across UMAPs (Extended Data Fig. 4). To predict layer-specific distributions of different neuronal types, we aligned the measured nuclei with recent datasets^16,17^, in which layer positions were validated experimentally (Fig. 1e, Extended Data Fig. 5,6).

To characterize schizophrenia-related differences in the DLPFC, we first examined whether the composition of the cortex was altered in schizophrenia patients. Analysis of normalized cell density on the joint UMAP embedding (Fig. 2a) and direct comparison of cell proportions (Fig. 2b, Extended Data Fig. 7a,b) showed a general decrease in GABAergic interneurons in schizophrenia (Extended Data Fig. 7a), affecting subtypes from all families, in particular those belonging to PVALB, SST and VIP (Fig. 2b, Extended Data Fig. 7b). This was countered by an increase in fraction of specific subtypes of principal neurons belonging to L2_3_CUX2_FREM3 and L4_5_FEZF2_LRRK1 (Fig. 2b, Extended Data Fig. 7a,b). As changes in proportion of one subtype could potentially skew the representation of other subtypes, we applied Compositional Data Analysis^18^ techniques to calculate robust estimates of compositional changes (see Methods). This analysis confirmed that schizophrenia was linked to a general increase in most subtypes of L2_3_CUX2 and L4_5_FEZF2 families of principal neurons, and a decrease in several interneuron subtypes, most notable for SST_CALB1 (Fig. 2c, Extended Data Fig. 7c).

**Figure 2.**
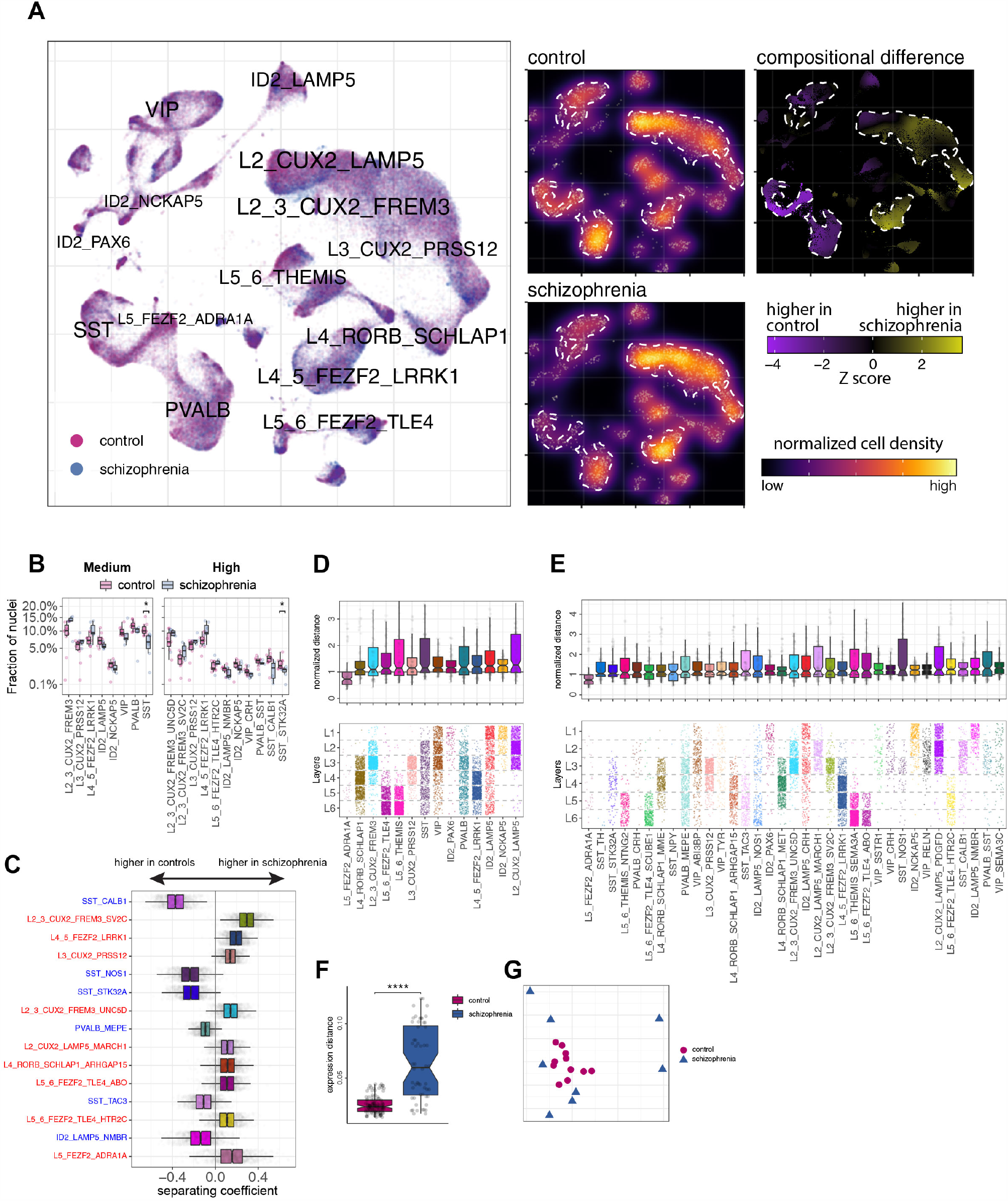
Compositional and transcriptomic changes in the cortex of patients with schizophrenia. A: Cell density differences between schizophrenia and control groups analyzed using UMAP embedding. (Left) Visualization of control and schizophrenia cells. (Middle) Cell density visualized from the control and schizophrenia samples, using UMAP embedding. (Right) Statistical assessment of the cell density differences. Student t-test was used, visualized as a Z score. B: Fraction of total nuclei represented by different subtypes are shown for schizophrenia and control groups. Normality was tested using Shapiro-Wilk test, equality of variances was tested using Levene’s test. As part of the data was not normally distributed Independent 2-group Mann-Whitney U test was used to test significance of differences for all group pairs. Multiple comparison correction was done using Benjamini & Hochberg method. p values: SST 0.01, SST_STK32A 0.042. Lower and upper hinges of the box correspond to the first and third quartiles. Line in the box corresponds to the median. Upper and lower whiskers correspond to the smallest and the largest values, and not more than 1.5 x inter-quartile range. C: Change in neuronal composition evaluated by Compositional Data Analysis. Top cell types distinguishing composition of control and schizophrenia samples are shown. The x axis indicates the separating coefficient for each cell type, with the positive values corresponding to neurons with increased abundance in schizophrenia, and negative to decreased abundance. The boxplots and individual data points show uncertainty based on bootstrap resampling of samples and cells. D,E: The boxplots showing the magnitude of transcriptional change between control and schizophrenia state for medium (D) and high (E) resolution annotations. The magnitude is assessed based on a Pearson linear correlation coefficient, normalized by the medium variation within control and schizophrenia groups (see Methods). The cell types are ordered based on the mean distance, with the most impacted cell types shown on the right. The lower panels show the predicted cortical layer positions. F: The boxplots showing inter-individual gene expression distances (based on Pearson correlation) within control and schizophrenia samples, averaged across all neuronal cell types. G: Multidimensional scaling visualization of the similarity of gene expression between all samples, based on the distances shown in F.

We then examined the extent to which the transcriptional state of different neurons is altered in schizophrenia. Using a simple expression distance measure based on the Pearson linear correlation, for each annotated type we compared the distances between schizophrenia and control samples with the average distance observed within schizophrenia and controls. At medium resolution, ID2 subtypes of GABAergic interneurons and L2_CUX_LAMP5 subfamily of principal neurons showed the largest difference between control and schizophrenia transcriptomes (Fig. 2d). When zooming in further, the majority of individual subtypes with the largest expression shifts belonged to GABAergic interneurons, such as VIP_SEMA3C, VIP_RELN, PVALB_SST, ID2_LAMP5_NMBR, ID2_NCKAP5 and SST_CALB1 (Fig. 2e). Interestingly, in terms of their layer positions most affected GABAergic interneuron subtypes were mainly confined to supragranular L2-3 (with some subtypes also being abundant in L1). This is particularly striking considering that most neurons of the SST and PVALB families of GABAergic interneurons reside primarily in deeper layers, whereas SST_CALB1 and PVALB_SST show high preference for supragranular layers (Fig. 1e).

Since the expression distance analysis also revealed substantial inter-patient variability, we examined the magnitude of transcriptional differences observed within the control and schizophrenia groups. We found that schizophrenia patients showed significantly greater inter-individual expression variability (Fig. 2f), a trend that was pronounced across all neuronal subtypes (Extended Data Fig. 7d,e). Consistently, when visualizing the distances between samples using multidimensional scaling, we found that while controls were grouped closer together, the schizophrenia samples were scattered in different directions (Fig. 2g). Since large within-group distances may occlude the extent to which different neuronal types are impacted by schizophrenia, we repeated the analysis of transcriptional shift magnitudes using a more complex distance measure that quantifies changes occurring only along the direction of the overall difference between schizophrenia and control states (see Methods). Specifically, for each cell type, we first determined a consensus pattern of expression differences between schizophrenia and controls. The distances between any pair of schizophrenia and control samples were then quantified by projecting the samples onto that consensus axis, and normalized by the distances expected from randomized assignment of samples to the control and schizophrenia groups. Consistent with the initial results, this analysis at medium resolution showed largest expression differences to be associated with L2_CUX_LAMP5 and subfamilies of GABAergic interneurons noted earlier (Extended Data Fig. 7f), and at high resolution the majority of most affected subtypes belonged to GABAergic interneurons of L2-3, such as PVALB_CRH, SST_CALB1, VIP_SEMA3C and ID2_LAMP5_NMBR (Extended Data Fig. 7g).

Overall, the strongest effect of schizophrenia on compositional and transcriptomic changes was attributed to subtypes of principal neurons and GABAergic interneurons in L2-3 of the DLPFC. Interestingly, such hotspot of changes in L2-3 subtypes might underlie the human-specific nature of schizophrenia, since there is a large body of evidence showing extensive evolution of L2-3 along the rodent-primate-human axis. Thus, L2-3 were shown to have the largest expansion out of all cortical layers in human relative to the rodent cortex^19^. Furthermore, based on unsupervised cluster analysis of neuronal morphology, the vast majority of L2-3 principal neurons are classified into human-specific clusters that are distinct from mouse and macaque neurons^20^. Finally, recent snRNA-seq studies showed high diversification of transcriptomic and morpho-electric properties of human L2-3 principal neurons^21^, and expansion of GABAergic interneurons residing in L2-3 in the PFC but not V1 in primates relative to mice^22^. As evolutionary divergence of interneurons is not well-understood, we quantified the inter-species expression distances based on a recent alignment of human, marmoset and mouse motor cortex^23^. The analysis showed ID2 and PVALB subfamilies positioned in the L2-3 region among the subtypes with the highest expression divergence in humans (Extended Data Fig. 8). Therefore, we focused on L2-3 DLPFC and to confirm the main effect of schizophrenia on L2-3 in the DLPFC, we performed immunohistochemical characterization of different families of GABAergic interneurons across all cortical layers in a set of 10 schizophrenia and 10 control brains that included those analyzed by snRNA-seq (Fig. 3a) (Supplementary Table 3). We labeled calretinin-expressing (CR+) GABAergic interneurons, the most enriched interneuron marker in human supragranular layers^24^. Based on our data they represent all VIP subtypes and two ID2 subtypes, ID2_NCKAP5 and ID2_PAX6 (*CALB2* gene codes for CR) (Fig. 1d). While the density of CR+ interneurons in L1 and L3-6 was similar between the conditions, there was a significant decrease specifically in L2 in schizophrenia (*p*<0.001, Fig. 3b,c, Supplementary Table 3). A conspicuous ‘L2 low CR phenotype’ was found in 50% of cases with schizophrenia.

**Figure 3.**
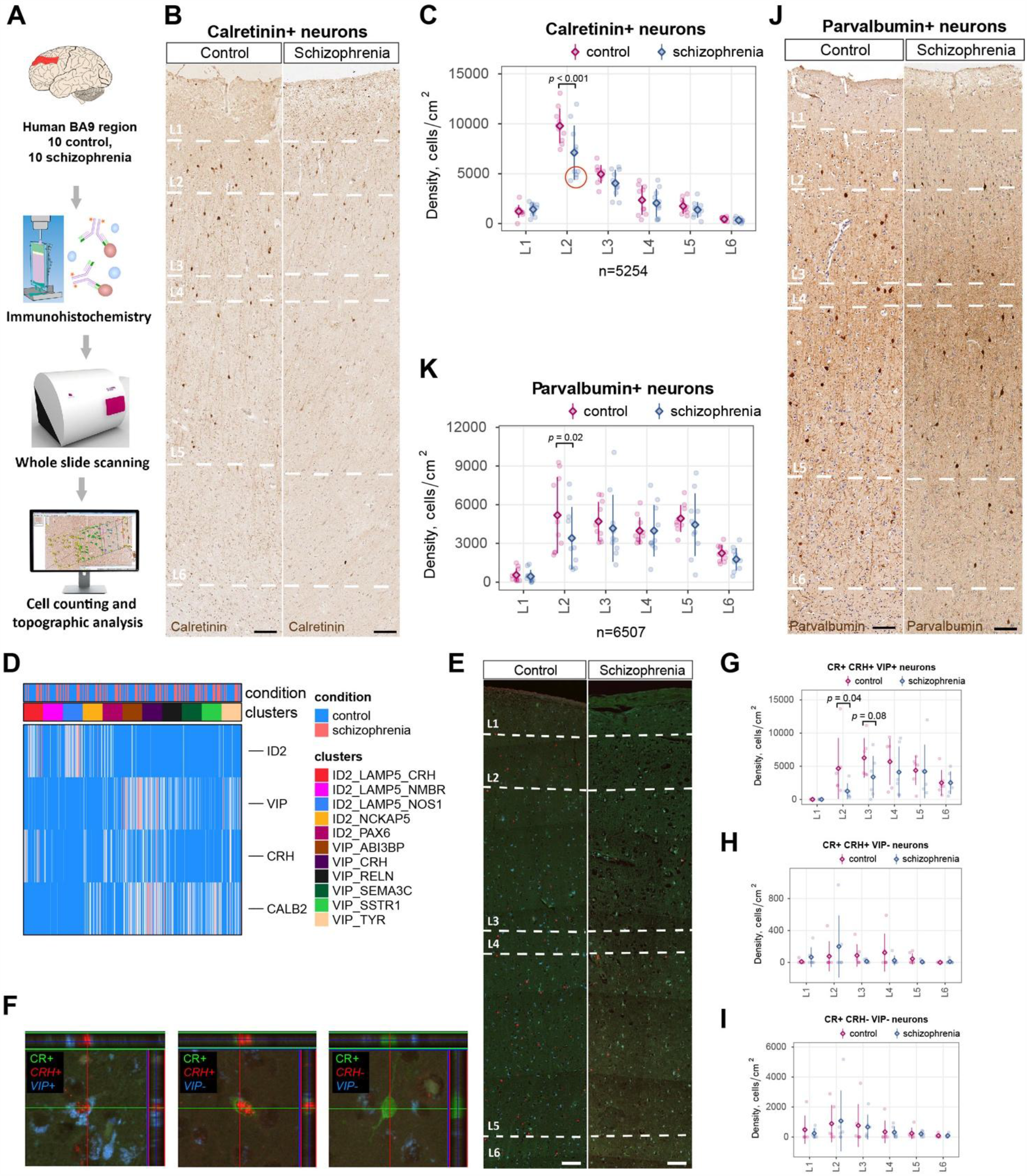
Changes in density and distribution of neuronal subtypes in the cortex of patients with schizophrenia analyzed by immunohistochemistry. A: Experimental scheme for immunohistochemical analysis. B,C: Representative images and layer-wise quantification of CR+ neurons in the DLPFC in patients with schizophrenia and control subjects; Linear Mixed Model analysis, scale bars: 100µm. D: Heatmap showing expression of major markers for subgrouping of ID2 and VIP subtypes of GABAergic interneurons. E,F: Overview of labeling with CR antibody, and smFISH probes for VIP and CRH mRNAs, and representative confocal images for three subgroups of ID2 and VIP subtypes that could be distinguished based on triple CR/VIP/CRH labeling in L2 DLPFC. G,H,I: Quantification of density for CR+/VIP+/CRH+ (VIP_CRH, VIP_ABI3BP, and VIP_TYR), CR+/VIP-/CRH+ (ID2_PAX6) and CR+/VIP-/CRH-(ID2_NCKAP5 and VIP_SSTR1) subgroups of ID2 and VIP subtypes, Linear Mixed Models analysis, scale bars: 100µm. J,K: Representative images and layer-wise quantification of PV+ neurons in the DLPFC in patients with schizophrenia and control subjects; Linear Mixed Models analysis, scale bars: 100µm. Diamonds on the plots represent mean, error bars +-SD.

To identify VIP and ID2 subtypes of GABAergic interneurons which might account for the observed CR+ density reduction in schizophrenia DLPFC, we implemented a set of the following markers – CR, VIP and CRH to further distinguish subpopulations of VIP and ID2 families. Thus, we measured the density of CR+/VIP+/CRH+, composed of VIP_CRH, VIP_ABI3BP, and VIP_TYR subtypes, CR+/VIP-/CRH+, labels for ID2_PAX6, and CR+/VIP-/CRH-, labels for ID2_NCKAP5 and VIP_SSTR1 (Fig. 3d-f). We confirmed that all of these subpopulations were mostly localized in L2-3, and observed a trend towards a reduced density for triple-positive CR+/VIP+/CRH+ subtypes of VIP interneurons in L2 (p=0.04) and in L3 (p=0.08) (Fig. 3g-i, Supplementary Table 3).

Parvalbumin expressing (PV+) interneurons (family PVALB) were also among those with the largest transcriptomic and compositional changes in our schizophrenia snRNA-seq data. Importantly, extensive data in animal models and patients establish a major role of PV+ interneurons in schizophrenia-related cognitive abnormalities^5,25,26^. We, therefore, analyzed the density of PV+ interneurons in the same cohort of DLPFC samples. There was a reduction in the density of PV+ interneurons in L2 in schizophrenia (*p*=0.02, Fig. 3j,k) (Supplementary Table 3), and such reduction was observed in 60% of cases with schizophrenia. Finally, to further validate snRNA-seq compositional findings for interneurons, we analyzed the distribution of calbindin (gene *CALB1*), which labels mainly SST_CALB1 neurons in L2-3 and detected a significant decrease in L2 density (p=0.007) in schizophrenia (Extended Data Fig. 9a,b, Supplementary Table 3). An additional decrease was also found in L4 for SST_NPY subtype (p=0.004) labeled by neuropeptide Y (NPY) (Extended Data Fig. 9c,d, Supplementary Table 3). Labeling for Nissl showed a decrease in total neuronal density in L2 (p=0.04) (Extended Data Fig. 9e,f), which should be associated with decrease in interneurons that was determined above, since labeling for principal neurons by SMI31.1 (the protein product of *NEFH* and *NEFM* genes) showed some visible increase in principal neurons in L2 in schizophrenia (Extended Data Fig. 9g,h). Overall immunohistochemical analysis of the DLPFC from schizophrenia patients confirmed compositional changes in supragranular layers revealed by snRNA-seq -a decrease in VIP/ID2 subtypes in L2, a decrease in PVALB and SST_CALB1 L2 subtypes, and a potential increase in principal neurons in L2.

To gain further insights into the transcriptomic changes in schizophrenia, we calculated differentially expressed genes (DEGs) for each subtype between schizophrenia and control samples. Interestingly, the subtypes with the highest number of DEGs or fraction of DEGs, estimated controlling for the differences in the number of the measured nuclei, reside preferentially in supragranular layers (Supplementary Table 4, Extended Data Fig. 10a,b). To explore pathways that might underlie schizophrenia-related changes in neuronal function, we extracted top up-/down-regulated genes for each subtype and calculated enrichment for Gene Ontology (GO) terms. The most significant GO terms downregulated were related to protein processing and transport, which was a common feature of subtypes in supragranular layers: L2_3_CUX2 principal neurons, VIP (VIP_SEMA3C, VIP_SSTR1), PVALB (PVALB_SST, PVALB_CRH) and ID2 (ID2_NCKAP5, ID2_PAX6) interneurons (Supplementary Table 5, Fig. 4a, Extended Data Fig. 10c). In contrast, the upregulated genes were enriched for functionally-relevant pathways in supragranular neuronal subtypes, which were related to synaptic plasticity, cell morphogenesis and development (Supplementary Table 5, Fig. 4b, Extended Data Fig. 10d). Downregulation of protein translation and upregulation of synaptic plasticity and morphogenesis might indicate developmentally-induced miswiring of supragranular neuronal circuits, alterations in neurotransmission and dysregulation of protein synthesis, assembly and transport in schizophrenia DLPFC. We noted that the upregulation of developmental/cell morphogenesis terms and downregulation of protein translation/processing terms similarly affected multiple subtypes of supragranular inhibitory neurons (Extended Data Fig. 11a,b). Clustering of GO terms by their enrichment level in each of the subtypes revealed groups of subtypes with similar enrichment patterns, which suggest them being in the same neuronal network. Thus, supragranular VIP_SEMA3C and ID2_NCKAP5 clustered with L3_CUX2_PRSS12, VIP_TYR with L2_3_CUX2_FREM3_SV2C, VIP_SSTR1 with PVALB_SST and ID2_PAX6, and PVALB_SST clustered together with L2_3_CUX2_LAMP5_PDGFD (Extended Data Fig. 11c,d). Additionally, we checked the expression of neurotransmitter receptors that have been proposed to contribute to schizophrenia-associated phenotypes or being targets of frontline anti-psychotic drugs^27,28^. Dopamine receptor *DRD2* was upregulated in VIP_CRH, GABA receptor *GABARA1* was decreased in ID2_PAX6, VIP_SSTR1 and PVALB_CRH. The kainate-type glutamate receptor *GRIK3* was upregulated in SST_CALB1 and SST_NOS1. The NMDR-type glutamate receptor *GRIN2A* was downregulated in ID2_LAMP5_NMBR and *GRIN2C* in PVALB_CRH, and decreased *OXTR* expression was detected in SST_NPY (Extended Data Fig. 11e). This analysis highlights receptor genes which were exclusively expressed in only one neuronal subtype, therefore may be used in future studies for selectively targeting neuronal subpopulations.

**Figure 4.**
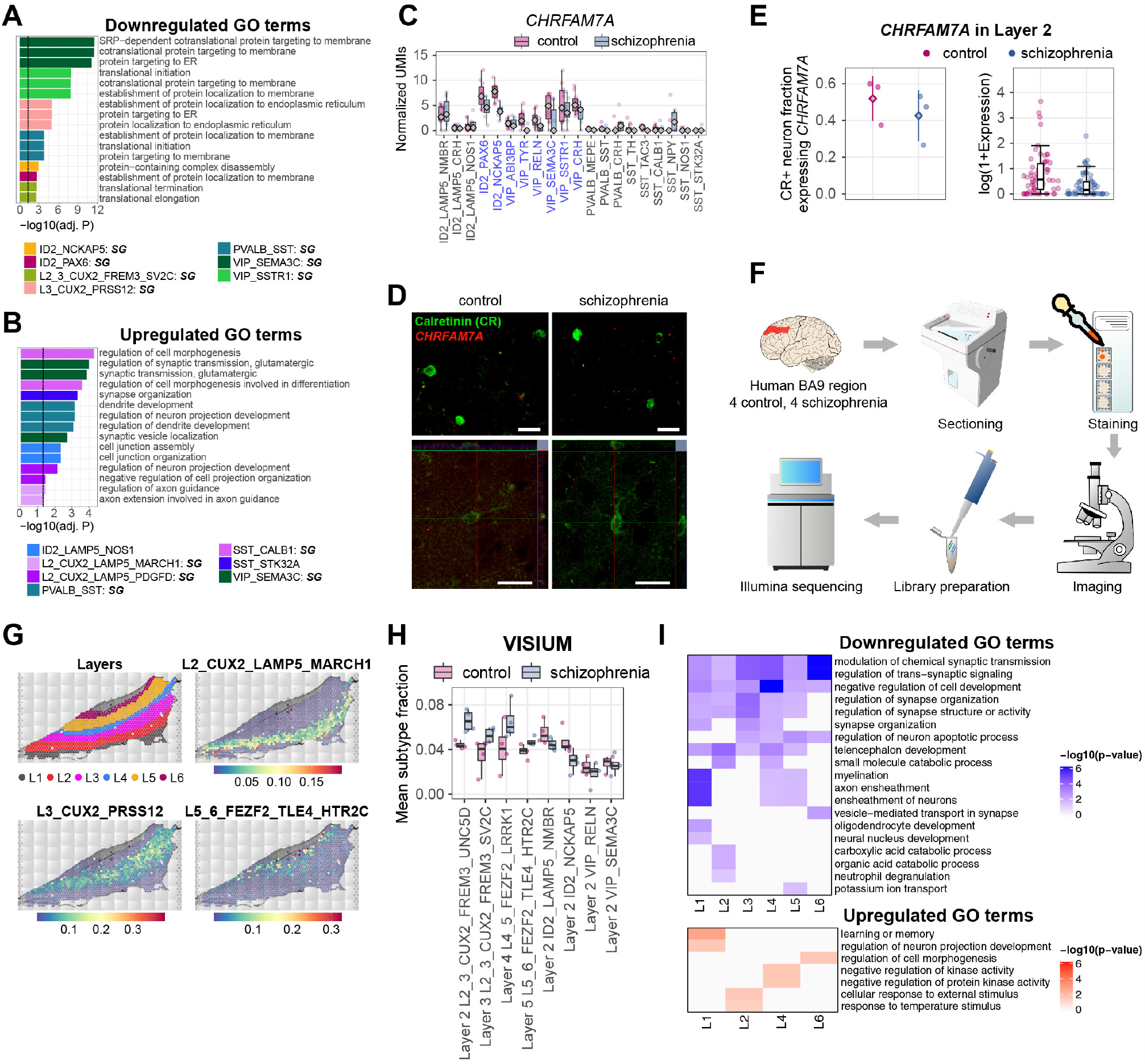
Schizophrenia-associated pathways in neuronal subtypes and spatial analysis of transcriptomic changes in the DLPFC in schizophrenia. A,B: GO terms enriched in genes downregulated (A) and upregulated (B) in the DLPFC neuronal subtypes in schizophrenia. SG -supragranular. C: Expression levels of *CHRFAM7A* across neuronal subtypes in the DLPFC identified by snRNA-seq. CR+ subtypes, highlighted in blue, show general downregulation of *CHRFAM7A* in schizophrenia. p values were estimated using differential gene expression Wald test in *DESeq2*, multiple comparison correction was done using Benjamini & Hochberg method. Expression levels were based on normalized pseudo-bulk mRNA UMI counts. Diamond represents median. D: Representative images for detection of CHRFAM7A mRNA in L2 CR+ neurons of patients with schizophrenia and control subjects. Scale bars: 20µm. E: Quantification of proportion of L2 CR+ neurons that co-localize with CHRFAM7A mRNA in SZ and control DLPFC (diamond - mean, error bars +-SD) and CHRFAM7A mRNA levels in L2 CR+ neurons. Expression values represent fraction of cell surface occupied by the mRNA of interest, values were ln(1 + expression) transformed for plotting. F: Experimental scheme for spatial transcriptomics measurements. G: Annotation of cortical layers in the DLPFC from spatial transcriptomics. H: Changes in neuronal composition in SZ DLPFC, identified in the spatial transcriptomics data. Normality was tested using Shapiro-Wilk test, equality of variances was tested using Levene’s test. As part of the data was not normally distributed Independent 2-group Mann-Whitney U test was used to test significance of differences for all group pairs. Multiple comparison correction was done using Benjamini & Hochberg method. I: GO terms enriched in the set of downregulated and upregulated genes in the neuronal subtypes in schizophrenia, observed in both snRNA-seq and spatial transcriptomics data. All boxplot panels: line inside the box represents median, lower and upper hinges of the box correspond to the first and third quartiles. Upper and lower whiskers correspond to the smallest and the largest values, and not more than 1.5 x inter-quartile range.

To validate DEGs in schizophrenia, we selected CR+ GABAergic interneurons (VIP and ID2 families) that showed one of the highest effects of schizophrenia in the above analyses and that are located mainly in L2-3. We confirmed again that most transcriptomic changes in CR+ neuronal subtypes were associated with downregulation of protein synthesis and metabolism, and upregulation of synaptic plasticity/morphogenesis/development (Extended Data Fig. 12a,b). From DEGs, we selected a gene coding for neurotransmission protein, *CHRFAM7A*, that was prominently expressed across CR+ neuronal subtypes and exhibited robust downregulation in schizophrenia (Fig. 4c). *CHRFAM7A* codes for a human-specific fusion protein that has acetylcholine receptor properties and was proposed to have a role in schizophrenia^29^. In a set of schizophrenia and control DLPFC sections (n=3 vs 3), we analyzed the expression of *CHRFAM7A* in CR+ cells and confirmed that the ratio of L2 CR+*CHRFAM7*+ to all L2 CR+ neurons was decreased by 18% in patients with schizophrenia. Furthermore, expression levels of *CHRFAM7* in L2 CR+ neurons were reduced in patients with schizophrenia by 77% (fig. 4d,e).

To enhance the spatial resolution and further validate schizophrenia-related compositional and transcriptomic differences in the DLPFC, we used a spatial transcriptomics assay (Fig. 4f). The spatial analysis confirmed our earlier estimates of layer-specific distribution of principal neurons and preferential localization of GABAergic interneurons (Fig. 4g, Extended Data Fig. 13). Importantly, we validated compositional changes in principal neurons that mainly affected supragranular layers (Fig. 4H, Extended Data Fig. 14 and 15a,b). We further performed gene ontology analysis of schizophrenia up- and down-regulated genes and found substantial overlap with the GO terms previously identified using snRNA-seq (Fig. 4i, Supplementary Table 6, 7). We identified a hotspot of transcriptomic changes in the upper layers of the cortex, consistent with our snRNA-seq results (Extended Data Fig. 15b,c). Nevertheless, we also noted transcriptional changes in the lower cortical layers and possible perturbations of glia giving us further insight into the high complexity of schizophrenia at the spatial tissue level.

We show that subtypes of GABAergic interneurons and principal neurons in supragranular layers have the largest compositional and transcriptomic changes in schizophrenia, and thus are likely to be the most vulnerable in schizophrenia. These findings should underline human-specific nature of schizophrenia as supragranular layers are the most expanded in humans^19^ and have increased their complexity morphologically, functionally, and transcriptomically^20,21^. Furthermore, such expansion has been proposed to contribute to complex cognitive functions in primates and in particular humans^30–32^. Accordingly, our data and earlier studies^7^ show that L2-3 in the human cortex have the highest complexity of GABAergic interneurons, since the number of GABAergic interneuron subtypes in each supragranular layer (L2 and L3) is higher than in any layer below (L4, L5, and L6). Importantly, the schizophrenia-associated differences do not belong to a selected family of neurons but involve multiple neuronal subtypes in L2-3, indicating a general network impairment. This widespread impact is also manifested by the orchestrated downregulation of protein processing and transport genes and upregulation of neuronal development/plasticity genes across multiple subtypes of principal neurons and GABAergic interneurons in supragranular layers. Similar changes have been previously noted in association with mental disorders based on genetics and bulk transcriptomics studies^10,33–35^.

Such a hotspot of schizophrenia-associated changes in the supragranular layers might arise during cortical development. Supragranular principal neurons in the DLPFC start intensive differentiation during the third trimester^36^. At the same developmental period, GABAergic interneurons that are destined for the supragranular layers arrive and spread^37^. Likewise, human cortical development at late gestation period is characterized by large-scale transcriptomic transition from embryonic to mature neuron transcriptome^34^, which includes re-shaping of transcriptome involved in neuronal differentiation and maturation^34,38^. Importantly, supragranular neurons have a highly protracted synaptic spine overproduction within cortico-cortical circuitries, where final synaptic removal coincides with appearance of schizophrenia symptoms^39^. Therefore, our data supports the theory that schizophrenia-associated developmental events occur in the DLPFC during late gestation due to disturbed arrival of GABAergic interneurons and differentiation of principal neurons.

Another conceptual finding in our work is that only some neuronal subtypes of the same family exhibit schizophrenia-associated changes, but not the whole family. For instance, PV interneuron family had been proposed to be impaired in schizophrenia before^5,25,26,40^, and here we show that only supragranular-enriched PVALB_SST and PVALB_CRH subtypes, but not the largest of PVALB subtypes, PVALB_MEPE, have extensive schizophrenia-associated transcriptomic changes. Same holds true for SST, where supragranular-enriched SST_CALB1 subtype showed by far the largest schizophrenia-associated effect compared to other SST subtypes, VIP (e.g. VIP_SEMA3C) and ID2 (e.g. ID2_NCKAP5) interneuron families.

Supragranular principal neurons were also proposed to be the most affected in a snRNA-seq study of another psychiatric disorder – autism spectrum disorder^41^. Thus, the resemblance of the effect of schizophrenia and autism on supragranular cortical layers may indicate a particular vulnerability of supragranular neuronal circuits to psychiatric disorders.

## DATA ACCESSIBILITY

The raw snRNA-seq data will be available at EGA at https://www.ebi.ac.uk/ena/submit/sra/#ega (due to data protection issues, we are in the process to secure the data protection agreement and the data will be available under “access by permission of data access committee” before the publication). Count matrices and code to reproduce the analysis will be available on github.

## ACKNOWLEDGEMENTS

We are grateful to Irina Korshunova, Maria Bako, Judit Kerti, Zsuzsa E Toth, Bozica Popovic and Maja Horvat for their technical assistance. We thank Carolyn Sloan, Hannah Brooks, Miklos Palkovits, Eva Renner, Michiel Kooreman and Debbie Lennard (Human Tissue Brain Bank at Semmelweis University, Oxford Brain Bank, the Netherlands Brain Bank, and Newcastle Brain Bank) for their kind assistance in procuring human brain tissue. We thank Zsolt Lang for his comments on statistical analysis of IHC and in situ hybridization data. We are also grateful to Alma Andersson for her comments on spatial transcriptomics analysis and Elena Ocheredko for initial help with data analysis. We thank BRIC’s single cell (Irina Korshunova), flow cytometry (Anna Fossum and Rajesh Somasundaram), imaging (Yasuko Antoku) and sequencing (Francisco Rodriguez Gonzalez) core facilities as well as CBMR sequencing core facility (Kristoffer Egerod) for their assistance during this work. The work was supported by Novo Nordisk Hallas-Møller Investigator grant (NNF16OC0019920), Lundbeck-NIH Brain Initiative grant (2017-2241), DFF-Forskningsprojekt1 (8020-00083B) to KK. The study was funded by the Institutional Excellence in Higher Education Grant (FIKP), STIA_18, TeT 2019-2021 and Departmental Start-up Grants (Semmelweis University) to IA. TT was supported by the ÚNKP-19-2 New National Excellence Programme of the Ministry for Innovation and Technology (Hungary). The work of DS, NH and ZP was supported by the Croatian Science Foundation grant no. 5943 (Microcircuitry of higher cognitive functions, PI: ZP), and by the European Union through the European Regional Development Fund, Operational Programme Competitiveness and Cohesion, grant agreement No. KK.01.1.1.01.0007, CoRE—Neuro.

## Supplementary Materials

Materials and Methods

Extended Data Fig 1-15

## Materials and Methods

### Human material

Tissue from subjects was provided by the Netherlands Brain Bank (NBB) Netherlands Institute for Neuroscience, Amsterdam (open access: www.brainbank.nl), the Oxford Brain Bank (OBB), the Human Brain Tissue Bank (HBTB, Semmelweis University, Hungary) and the Newcastle Brain Bank, UK (Newcastle). All material was collected from donors from whom written informed consent had been obtained by the NBB, OBB, HBTB or Newcastle for brain autopsy and use of material and clinical information for research purposes. Importantly, to form the sample set for our study, we tried to select samples to avoid large variability in age, post-mortem interval and other major confounders, which could increase the noise and mask biologically-relevant data. The demographic characteristics of the cohort are shown in Supplementary Table 1 and 3. The mean values of age, gender, post-mortem interval and time in paraformaldehyde were not significantly different in the diagnostic groups examined (Extended Data Fig. 1). Nevertheless, the effect of medication and lifestyle cannot be excluded, and the effect on cell types that we observe could be in part due to particular lifestyle of patients with schizophrenia or medication that they use. Sampling was done by assistants of the NBB, OBB, HBTB and Newcastle supervised by trained neuropathologists. Both paraffin-embedded sections and fresh-frozen tissue blocks were provided from the dorsolateral prefrontal cortex (Brodmann area 9).

The samples that were used for each type of the analysis are indicated in Supplementary Table 3 (see “Type of analysis” column). Although most samples were used for both snRNA-seq and immunohistochemistry, few samples were studied only by one of these methods, due to the limited availability of tissue in brain banks or due to availability of only fresh-frozen or only formalin-fixed tissue. All samples for spatial transcriptomics were also analyzed by snRNA-seq.

### Nuclei isolation

Wherever possible schizophrenia and control samples were processed in parallel, 2 samples per experiment. Tissue was removed from −80 °C and placed in 1ml Dounce homogenizer with 1 ml of chilled Homogenization buffer (250 mM Sucrose, 25 mM KCl, 5 mM MgCl2, 10 mM Tris pH 8.0, 1 mM DTT, 1x protease inhibitor (Roche, 11873580001), 0.4 U/ul RNAse inhibitor (Takara, 2313B), 0.2 U/ul Superasin (Invitrogen, AM2696), 0.1% Triton X-100) on ice. Tissue was homogenized with 5 strokes of loose pestle and 15 strokes of tight pestle. Homogenate was filtered through a 40 um cell strainer and spun down at 1000 g for 8 min at 4 °C. The pellet was gently resuspended in a Homogenization buffer with the final volume of 250 ul on ice. Suspension was mixed with 250 ul of 50% Iodixanol solution (25 mM KCl, 5 mM MgCl2, 10 mM Tris pH 8.0, 50% Iodixanol (60% stock from STEMCELL Technologies, 7820), 1x protease inhibitor (Roche, 11873580001), 0.4 U/ul RNAse inhibitor (Takara, 2313B), 0.2 U/ul Superasin (Invitrogen, AM2696), 1 mM DTT) and overlaid on top of 29% Iodixanol solution (25 mM KCl, 5 mM MgCl2, 10 mM Tris pH 8.0, 29% Iodixanol, 1x protease inhibitor (Roche, 11873580001), 0.4 U/ul RNAse inhibitor (Takara, 2313B), 0.2 U/ul Superasin (Invitrogen, AM2696), 1 mM DTT) in the ultracentrifuge tube (Beckman Coulter, 343778) on ice. Gradients were spun down in the ultracentrifuge (Beckman Coulter, MAX-XP) using a swing bucket rotor (Beckman Coulter, TLS 55) at 14000 gmax for 22 min at 4 °C with slow acceleration and deceleration. Following, supernatant was carefully removed and pellets were resuspended in ice-cold BSA solution (1x PBS (AM9625, Ambion), 0.5% BSA (VWR, 0332-25G), 1 mM DTT, 2.4 mM MgCl_2_, 0.2 U/ul RNAse inhibitor (Takara, 2313B)) and incubated on ice for 15 min for blocking. Suspension was split into experimental and FACS control (isotype control, negative control, 7AAD only control, NeuN only control) samples. Experimental sample and NeuN only control were stained with antibody against the neuronal marker NeuN (Millipore, MAB3777x, 1ug/ul) at 1:1890 dilution for 10 min at 4 °C protected from light. Isotype control was stained using control antibody (STEMCELL Technologies, 60070AD.1, 0.2 ug/ul) at 1:378 dilution for 10 min at 4 °C protected from light. Following, 1 ml of BSA solution was added, suspensions were mixed and centrifuged for 10 min at 1000g at 4 °C using a swing bucket rotor. Pellets were resuspended and filtered through 35 um strainers. Experimental sample, isotype control and 7AAD only control were supplemented with 0.75 ul of 7AAD (1 mg/ml) per 0.5 ml of suspension. FACS was done using BD FACSAria III sorter using 75 um nozzle and controlled by BD FACSDiva 8.0.1 software. Compensations were done based on single color controls. Gates were set based on FACS controls. Nuclei were selected based on FSC-A/SSC-A gate, doublets removed using FSC-W/FSC-H and SSC-W/SSC-H gates, nuclei were further selected based on 7AAD staining and neuronal nuclei were sorted based on NeuN staining. Nuclei were kept at 4 °C during FACS and sorted into 5 ul of BSA solution at 4 °C. Nuclei were processed directly for 10x library preparation.

Representative FACS plots on the Extended Data Fig. 2 were prepared using FCS Express 7 Plus v7.04.0014 (De Novo Software).

### 10x library preparation and sequencing

Chromium Single Cell 3, Reagent Kits v3 (10x Genomics, PN-1000075) were used for library preparation according to a standard protocol. In brief, nuclei were counted under microscope, mixed with reverse transcription mix, and partitioned together with v3 Gel Beads on Chromium Chip B (10x Genomics, PN-1000073) into GEMs using Chromium Controller (10x Genomics, PN-120223). Following reverse transcription samples were frozen at −20 °C. Within a week samples from several 10x runs were processed together for cDNA clean-up and pre-amplification (12 PCR cycles). After SPRIselect clean-up cDNA was quantified and frozen at −20 °C. In general, the same quantity of cDNA was used during fragmentation, end-repair and A-tailing for most samples. Following, fragments were cleaned-up using SPRIselect reagent and processed through steps of adapter ligation, SPRIselect clean-up, and sample index PCR (using Chromium i7 Sample Indices (10x Genomics, PN-120262) for 11 PCR cycles). Following, libraries were cleaned-up with SPRIselect reagent, quantified using Qubit HS dsDNA Assay Kit (Thermo Fisher Scientific, Q32854) and Qubit Fluorometer and also using High Sensitivity DNA Kit (Agilent, 5067-4626) and Agilent 2100 Bioanalyzer. Libraries were pooled according to the expected amount of nuclei per sample; pool was quantified and sequenced using two NovaSeq 6000 S2 (Illumina, 20012861) runs on Illumina NovaSeq 6000 System (Illumina 20012850) controlled by NovaSeq Control Software v1.6.0. Libraries were sequenced using 28 cycles for read 1, 8 cycles for i7 Index and 94 cycles for read 2. On average, there were xxx reads and 4,413 genes per nucleus.

### Single nuclei sequencing data analysis

Quality of sequencing data was checked using Illumina’s *interop* v1.1.10 tool. Primary data analysis was performed using cellranger v3.1.0 (10x Genomics). Release 97 of human genome from Ensembl was used:

http://ftp.ensembl.org/pub/release-97/fasta/homo_sapiens/dna/Homo_sapiens.GRCh38.dna.primary_assembly.fa.gz

http://ftp.ensembl.org/pub/release-97/gtf/homo_sapiens/Homo_sapiens.GRCh38.97.gtf.gz

GTF file was modified to create pre-mRNA reference (“transcript” were renamed to “exon” and retained in the final gtf: *awk ‘BEGIN{FS=“\t”; OFS=“\t”} $3 == “transcript”{$3=“exon”; print}’ Homo_sapiens*.*GRCh38*.*97*.*gtf > Homo_sapiens*.*GRCh38*.*97*.*PREmRNA*.*gtf*). Then, the reference genome was created using *cellranger mkref*. Illumina .bcl files were de-multiplexed using *cellranger mkfastq* pipeline; reads were mapped against reference genome and UMIs were counted using *cellranger count* pipeline. Filtered feature barcode matrices were used for further secondary analysis.

Secondary analysis was performed using pagoda2 (https://github.com/kharchenkolab/pagoda2) and Conos (https://github.com/kharchenkolab/conos). To eliminate potential doublets, scrublet scores were determined for each dataset, and only cells with the score below 0.3 were considered for downstream analysis. Each individual dataset was normalized using pagoda2 with default parameters, requiring at least 500 molecules per cell. Different samples were then aligned using Conos with default parameter settings (PCA space with 30 components, angular distance, mNN matching, k=15, k.self=5), and UMAP embedding was estimated using default parameter settings.

Normality of distribution of fractions of nuclei derived from different neuronal subtypes (Extended Data Fig. 2, 7) was tested using Shapiro-Wilk test (*shapiro*.*test*), equality of variances were tested using Levene’s test (*leveneTest*). As part of the data was not normally distributed Independent 2-group Mann-Whitney U test (*wilcox*.*test*) was used to test significance of differences for all group pairs. Multiple comparison correction was done using Benjamini & Hochberg method.

Gene expression plots (Fig. 4c and Extended Data Fig. 11e) were based on normalized pseudo-bulk gene expression levels. UMI counts were summed up separately for each gene across all cells derived from a specific subtype-sample combination. Subtype-sample combinations with less than 10 cells were filtered out. Pseudo-bulk counts were then normalized: divided by the sum of all counts across all genes in any particular subtype-sample combination and multiplied by 10^6. p values were estimated using differential gene expression Wald test in *DESeq2*, multiple comparison correction was done using Benjamini & Hochberg method.

### Estimation of differential cell density

To estimate differential cell density between sample groups, we first compute kernel density in joint embedding space for each sample using ks R package (bin=400). Then quantile normalization was used to normalize the density matrix across samples. Average density of each sample group was shown in Fig. 2a. To impute the differential cell density between sample groups, we performed t-test between sample groups in each girded bin. To avoid noise from the background, we filter bins with at least 1 cell and the Z score is shown as heatmap.

### Expression distance analysis

Expression differences between matching subpopulations were determined by first estimating “mini-bulk” (or meta-cell) RNA-seq measurements for each subpopulation in each sample. Briefly, in each dataset, the molecules from all cells belonging to a given subpopulation were summed for each gene (i.e. forgetting cellular barcodes). The distance between the resulting high-coverage RNA-seq vectors was calculated using Pearson’s linear correlation on log transformed values. To attenuate the impact of the differences in the number of cells, cell types with more than 500 cells were randomly downsampled to 500 cells. A total of 100 sampling rounds were performed. To obtain final normalized distance estimates (Fig. 2d,e), the expression distances of sample pairs between the conditions (control and schizophrenia) were normalized by the median expression distance of pairs within the control and schizophrenia conditions.

Overall expression distances between samples (Fig. 2f) were determined as a normalized weighted sum of correlation distances across all cell subpopulations contained in both samples, with the weight equal to the subpopulation proportion (measured as a minimal proportion that the given cell subpopulation represents among the two samples being compared). Expression distances between samples are projected to 2D space using MSD (Fig. 2g). To illustrate inter-individual expression distance within each sample group, Inter-sample expression distance is shown as boxplot (Fig. 2f).

To determine expression distance along a consensus direction (Extended Data Fig. 7f,g), for each cell type a consensus expression shift between control and schizophrenia conditions was determined by calculating a trimmed mean (trim=0.1) log2 fold expression change for each gene. The resulting shift vector was normalized by its Frobenius norm to obtain the consensus shift vector *v*. For each pair of samples, the final distance was determined as a dot product between the log2 fold expression change vector for a given pari, and the normalized consensus shift vector *v*.

### Estimation of expression shifts across species

To estimate the extent of expression shifts across species, we integrated schizophrenia data with a dataset from Allen Institute that integrate single-cell transcriptomic datasets in primary motor cortex across human, mouse and marmoset^1^. Cell labels were mapped to schizophrenia cell types using label transfer function based on the joint embedding of integrated clusters. For each of the major cell types, the proportion of expression shift was estimated by calculating the correlation metric between human/marmoset and human/mouse. The cells in each major cell type were randomly subsampled to 500 and the entire process was repeated by 20 rounds.

### Compositional analyses

We performed compositional analyses as described elsewhere^2^. In short, isometric log-ratio transformations were applied to cell type fractions followed by canonical discriminant analyses using the *candisc* package to obtain weighted contrasts between cell types in schizophrenia and control samples. Random cell subsamlings were applied to evaluate robustness and statistical significance of the separating coefficients. In total, 1,000 subsamplings were performed each time evaluating 1,000 randomly sampled cells from both groups. The separating coefficients are plotted.

### Differentially expressed genes

Differential expression (DE) was performed on pseudobulk gene counts using the Wald test in *DESeq2*^3^ through the getPerCellTypeDE() function in *Conos*^4^. In short, gene counts per cell type were collapsed per sample to create pseudobulk gene counts. Benchmarking^5^ has proved this to be an effective approach for DE gene discovery in single-cell studies. Then, DE genes are determined using a grouping factor as a model for expression (schizophrenia versus control). DE results were used as input for gene ontology (GO) estimations.

### Gene ontology

GO enrichment was investigated using the *clusterProfiler*^6^ package. Enrichment was evaluated per cell type for GO BP categories on sets of top 300 up- and down-regulated genes separately based Z score. All expressed genes were used as background. Enriched BP categories with p-value < 0.05 were clustered into 20 clusters based on the similarity of the genes involved in the categories using binary distance measures. The clusters were named based on the most significantly enriched category.

### Estimation of cortical layer positions of our neural cell subtypes and their correspondence to the Allen institute neural subtype annotations

Locations of neural cell subtypes were estimated from the Allen Institute human motor cortex (M1C), temporal cortex (MTG) and anterior cingulate cortex (CgG) datasets^7,8^:

https://transcriptomic-viewer-downloads.s3-us-west-2.amazonaws.com/human/transcriptome.zip

https://transcriptomic-viewer-downloads.s3-us-west-2.amazonaws.com/human/sample-annotations.zip

To minimize differences between our (KU) and Allen Institute single nuclei datasets we re-mapped KU samples against Allen Institute genome reference:

RefSeq Genomic FASTA https://www.ncbi.nlm.nih.gov/assembly/GCF_000001405.28/

GTF http://celltypes.brain-map.org/api/v2/well_known_file_download/502175284

Allen Institute GTF file was modified using gffread^9^ as follows: *gffread rsem_GRCh38*.*p2*.*gtf -T -v -F --keep-genes --keep-exon-attrs -o rsem_GRCh38*.*p2_2*.*gtf*

Then, gene_symbol were renamed into gene_name:

*sed ‘s/gene_symbol/gene_name/g’ rsem_GRCh38*.*p2_2*.*gtf > rsem_GRCh38*.*p2_3*.*GTF*

pre-mRNA GTF was created as follows:

*awk ‘BEGIN{FS=“\t”; OFS=“\t”} $3 == “transcript” {$3=“exon”; print}’ rsem_GRCh38*.*p2_3*.*GTF > rsem_GRCh38*.*p2_3_premRNA*.*GTF*

Following, Allen Institute genome reference was created using *cellranger mkref*. KU MB14, MB15, MB17 sample reads were re-mapped against the Allen Institute reference genome and UMIs were counted using the *cellranger count* pipeline. Pagoda2 v0.1.1 run under R version 3.5.3 in RStudio version 1.2.1335 was used to create pagoda objects from feature barcode matrices using *basicP2proc* with n.odgenes = 3000, min.cells.per.gene = 0, min.transcripts.per.cell = 0 parameters. Conos v1.2.1 was used to create conos objects with *Conos$new*. KU single nuclei and Allen Institute datasets (either MTG, M1C or CgG) were integrated using *con$buildGraph* with parameters *k = 30, k*.*self = 5, space = “PCA”, ncomps = 30, n*.*odgenes = 3000, matching*.*method = “mNN”, metric = “angular”*; parameters *alignment*.*strength* and *same*.*factor*.*downweight* were individually tweaked to optimize integration between datasets.

Following, high or medium resolution subtype annotations were transferred from KU single nuclei to Allen Institute datasets using *conos$propagateLabels*. Subtype labels were given based on the highest probability of nucleus to belong to a certain subtype. Then, cortical layer positions of Allen Institute nuclei (derived from microdissections of cortical layers before FACS sorting of nuclei) were used for plotting cortical positions of KU neural cell subtypes (Fig. 1 and Extended Data Fig. 6). Correspondence map between Allen Institute and KU annotation of neural cell subtypes (Extended Data Fig. 5) was based on the same transferred annotations. Combinations of KU subtype – Allen Institute subtype labels found in ≤ 7% nuclei from a particular KU subtype were filtered out. Number of nuclei on both KU and Allen axes on Extended Data Fig. 5 were square root transformed twice to increase visibility of rare subtypes (SST_NPY, SST_TH) on the map.

### Human cortex slice preparation for Visium analysis

Tissue slices were prepared according to the standard Visium tissue preparation guide (10x Genomics, CG000240). Slices were mounted on Visium Gene Expression Slides (10x Genomics, 2000233) from Visium Spatial Gene Expression Slide Kit (10x Genomics, PN-1000185), two from patients with schizophrenia and two from controls on the same slide.

### Tissue staining, imaging and Visium library preparation

Tissue slices were processed according to the standard Visium Methanol Fixation and H&E Staining guide (10x Genomics, CG000160). In brief, slices were fixed in methanol, stained with Hematoxylin followed by bluing and Eosin staining. Slices were covered with mounting medium (85% Glycerol, 4.4 U/ul of RNAse Inhibitor (Takara, 2313B), 1 mM DTT) and coverslips before imaging. Slides were imaged using Axio Observer.Z1/7 confocal microscope (Zeiss) controlled by Zen 2.3 software. Images were pre-processed using FIJI 1.52p^10^.

Straight after imaging coverslips were removed using 3x SSC Buffer and samples were directly processed according to the standard Visium Spatial Gene Expression protocol (10x Genomics, CG000239) using Visium Spatial Gene Expression Slide & Reagent Kit (10x Genomics, PN-1000184). Tissue was permeabilized during 12 min, and mRNA was reverse transcribed, followed by second strand synthesis and denaturation. cDNA was quantified using KAPA SYBR FAST qPCR Master Mix (Roche, KK4600) on LightCycler 480 Real-Time PCR System (Roche) to determine an optimal amount of PCR cycles required for cDNA preamplification. 17 cycles were used for cDNA pre-amplification followed by cDNA clean-up with SPRIselect reagent (B23318, Beckman Coulter) and cDNA quantification. Same amount of cDNA was used for fragmentation, end repair and A-tailing step. Following, SPRIselect clean-up, adaptor ligation, another SPRIselect clean-up and sample index PCR (13 cycles, using Dual Index Kit TT (10x Genomics, 1000215)) steps were performed. Libraries were cleaned-up using SPRIselect reagent, quantified using High Sensitivity DNA Kit run on Agilent 2100 Bioanalyzer and also KAPA Illumina library quantification kit (Roche, 7960140001) run on LightCycler 480. Tissue-covered spots were quantified using Loupe Browser v4.1.0 (10x Genomics) and libraries were pooled according to their concentration and spot occupation on slides. Library pool was quantified on Bioanalyzer and with qPCR and sequenced using one NextSeq 500/550 High Output Kit v2.5 (Illumina, 20024907) on Illumina NextSeq500 using these parameters: read 1 28 cycles, i7 index 10 cycles, i5 index 10 cycles, read 2 90 cycles.

### Visium spatial transcriptomics data analysis

Primary data analysis was performed using spaceranger v1.1.0 (10x Genomics). See metadata in Supplementary Table 8. Two reference genomes were used. Release 97 of human genome from Ensembl was used to generate feature barcode matrices for differential gene expression and GO analyses:

http://ftp.ensembl.org/pub/release-97/fasta/homo_sapiens/dna/Homo_sapiens.GRCh38.dna.primary_assembly.fa.gz

http://ftp.ensembl.org/pub/release-97/gtf/homo_sapiens/Homo_sapiens.GRCh38.97.gtf.gz

While Allen Institute reference was used to prepare feature barcode matrices for Stereoscope^10^ cell type deconvolution:

RefSeq Genomic FASTA

https://www.ncbi.nlm.nih.gov/assembly/GCF_000001405.28/

GTF file

http://celltypes.brain-map.org/api/v2/well_known_file_download/502175284 Allen Institute GTF file was modified to rename gene_symbol into gene_name:

*sed ‘s/gene_symbol/gene_name/g’ rsem_GRCh38*.*p2*.*gtf > rsem_GRCh38*.*p2_GENE_NAME*.*GTF*

Following, reference genomes were created using *spaceranger mkref*. Illumina .bcl files were de-multiplexed using *spaceranger mkfastq* pipeline; reads were mapped against reference genomes and UMIs were counted using *spaceranger count* pipeline. Filtered feature barcode matrices were used for further analysis. Additionally, low quality tissue areas were excluded manually and cortical layers were annotated by the experienced histologist using Loupe Browser v4.1.0 (10x Genomics). White matter was excluded from analysis based on manual cortical layer annotation.

To investigate cell type composition of the spatial transcriptomics samples we performed deconvolution using Stereoscope. Because the 10x snRNA-seq samples, produced for this study, were depleted from non-neural cells during FACS, we added non-neural cells from Allen Institute Cell Types Database: RNA-Seq Data Human M1 - 10x genomics (2020)^11^ to our neuronal nuclei: https://idk-etl-prod-download-bucket.s3.amazonaws.com/aibs_human_m1_10x/matrix.csv https://idk-etl-prod-download-bucket.s3.amazonaws.com/aibs_human_m1_10x/metadata.csv Non-neural nuclei were selected based on class_label “Non-neural” in the metadata. Medium resolution annotation of Allen institute non-neural nuclei was derived from subclass_label in the metadata, and high resolution annotation was derived from cell_type_alias_label in the metadata. We used Seurat v3.1.4 on the MB11 spatial sample to find 3000 highly variable genes, which were later used for the deconvolution. For the deconvolution, we applied Stereoscope using the merged scRNA-seq count matrix to all spatial samples for different resolution of annotation. For medium resolution we used 10000 epochs (“*-sce*” parameter) and batch size of 1000 (“*-scb*”) for fitting scRNA-seq data and 20000 epochs (“*-ste*”) with batch size of 1000 (“*-stb*”) for fitting spatial data. And for high resolution we used *sce*=20000, *scb*=1000, *ste*=30000 and *stb*=1000.

Fractions of deconvoluted cell subtypes per Visium spot were plotted on top of histological images (Fig.4g and Extended Data Fig. 13) using Seurat v3.2.0^12^ run in R. Seurat objects were created using *Load10X_Spatial*. Manual layer annotation metadata was added to the objects; Stereoscope deconvolution predictions were added to the Seurat objects using *CreateAssayObject*. Plotting was performed using *SpatialPlot*, high resolution subtypes are shown for the MB11 sample.

Stereoscope predicted fractions of subtypes were averaged per sample per subtype (Fig. 4h and Extended Data Fig. 15a) for high and medium annotations; or averaged per sample per layer (Fig. 4H and Extended Data Fig. 14) per subtype. Normality of distribution was tested using Shapiro-Wilk test (*shapiro*.*test*), equality of variances was tested using Levene’s test (*leveneTest*). As part of the data was not normally distributed Independent 2-group Mann-Whitney U test (*wilcox*.*test*) was used to test significance of differences for all group pairs. Multiple comparison correction was done using Benjamini & Hochberg method.

Given the manual layer annotation, differentially expressed genes were estimated using getPerCellTypeDE function from the Conos package ^4^. Then, Gene Ontology terms (GOs) were estimated for each layer using 500 genes with the highest / lowest Z-score for up- or down-regulated GOs correspondingly using the clusterProfiler package ^6^. To group GOs that were enriched based on the same genes, we clustered them using Jaccard similarity of the gene sets. For that, pairwise Jaccard distances were estimated on the sets of enriched genes for all GOs with adjusted p-values below 0.05. Then, hierarchical clustering was performed on these distances (hclust function with method=‘ward.D2’) and the hierarchy was trimmed to 20 clusters. The p-value of the resulting cluster for a layer was estimated as minimum across all p-values of the cluster’s GO terms. The name of a cluster was set to the name of its term with the lowest geometric mean of p-values across all layers. To compare results of GO analysis on spatial data to the GOs discovered on snRNA-seq data, we used only spatial GO that were significantly enriched in snRNA-seq data, adjusted p-values only on them using Benjamini-Hochberg correction (which allowed to increase power of the test) and visualized the filtered spatial GOs with adjusted p-value below 0.05.

### Immunohistochemistry

Consecutive paraffin embedded brain sections were stained for the 5 antigens investigated and for Nissl. One section per subject was used per immunostaining (Supplementary Table 3). Immunohistochemical analysis was done as described in earlier studies^13^. Briefly, the sections were dewaxed through a graded alcohol series and treated with 3% H_2_O_2_ solution (in phosphate buffered saline, pH 7.4) for 30 minutes. Antigen retrieval was applied by autoclaving the slides in citrate buffer (0.01 M, pH 6.0) at 121 °C for 10 minutes. The following primary antibodies were used: anti-calretinin (rabbit, 1:300, Chemicon, AB5054), anti-neuropeptide Y (rabbit, 1:250, Abcam, ab30914), anti-calbindin (mouse, 1:300, Swant, D28k-300), anti-parvalbumin (rabbit, 1:500, Abcam, ab11427) and anti-SMI31.1 (mouse, 1:500, Biolegend, 837801) in Tris-buffered saline/Triton TMX-100 (pH 7.4) for 1 hour (100 µl/section). Sections were then incubated with horseradish peroxidase-linked secondary antibody from the Envision Kit (Dako, K-5007) for 1 hour (100 µl/section) and labelling was visualized by DAB from the same Envision Kit applied for 90 seconds (100 µl/section). During incubation with primary and secondary antibodies slides were put into Sequenza System coverplates and rack (Thermo Scientific, 72110017, 73310017). Two rinses with Tris-buffered saline/Triton TMX-100 (pH 7.4) were applied between the above-described steps of immunohistochemistry (1000 µl each). Haematoxylin nuclear counterstain was applied for 20 seconds. Sections were dehydrated through a graded alcohol series and coverslipped with DePeX. No labelling was observed when primary antibodies were omitted from the protocol.

### Image analysis and quantification

Sections were digitized using a slidescanner (3D Histech) at 40x magnification. The regions of interest (2mm x 1mm columns with all cortical layers) were outlined using the ImageScope programme (Aperio, v11.2.0.780). Designation of cortical layers (BA9) were outlined based on Nissl and SMI31.1 stainings in good agreement with^14^. Cross-sectional areas analysed in this study are shown in Supplementary Table 3. The longest diameter of every CR-, PV-, CB-, SMI31.1- and NPY-ip cell body in the region of interests were manually measured as described in^13^. Three investigators contributed to the quantification and all were blinded to the diagnoses of the subjects through random coding of the subject identifiers. Neuronal cell bodies with a diameter >4 µm and a width >2 µm were included in further statistical analysis.

### Single molecule fluorescent in situ hybridization (smFISH)

Sections from schizophrenia and control cases were processed for simultaneous detection of target mRNAs (CHRFAM7A, CRH, VIP) in combination with calretinin immunohistochemistry as previously described ^15,16^. Paraffin embedded sections were first deparaffinized and treated with 3% H2O2 for 10 minutes, rinsed in 0.01M phosphate-buffered saline (PBS, pH 7.4). Sections were then processed for fluorescent in situ hybridization (RNAscope) using RNAscope Multiplex Fluorescent Kit V2 (Advanced Cell Diagnostics, Inc., Newark, CA, USA; cat 323110) according to the manufacter’s protocol. Briefly, sections were treated with 100% ethanol, incubated in antigen retrieval buffer maintained at a boiling temperature for 10 minutes, rinsed in PBS, and immediately treated with protease plus for 30 minutes at 40°C. RNAscope® Probe – Hs (homo sapiens)-CHRFAM7A-C1 (cat 833991) or CRH-C1 (cat 475211) for detection of target mRNAs and RNAscope® Probe Hs-VIP-C2 (cat 452751-C2) for the detection of VIP mRNA were hybridized for 2h at 40°C. After hybridization, sections were processed to amplification of target probes and then visualized using the Tyramide Signal Amplification (TSA™) (Perkin Elmer, Waltham, MA, USA [Wang et al., 2012]), fluorescein for CHRFAM7A or CRH mRNA, and Cy3 for VIP mRNA. Following the RNAscope assay, sections were rinsed in PBS and processed for immunohistochemical detection of calretinin. Sections were incubated for 1 hour at room temperature in 3% normal donkey serum (NDS, Vector Laboratories, Inc., Burlingame, CA, USA) and overnight at 4°C in the rabbit polyclonal antibody directed against calretinin (1:300, Millipore, Burlinghton, MA, USA; AB5054) diluted in PBS containing 0.3% Triton X-100 and 1% NDS. After 3 washes in PBS, the sections were incubated for 2 hours in Alexa 647-conjugated donkey anti-rabbit (1:2000; ThermoFisher), 3 times washed in PBS and mounted by Vectashield (Vector Laboratories, Inc., Burlingame, CA, USA).

### Statistical analysis of immunohistochemistry and smFISH data

Results are presented as means ± standard deviation. Student’s t-test (unpaired, two-tailed) was used to assess if the means of variables (such as age, PMI, cortical width, neuronal diameters) between the two diagnostic groups were significantly different (Supplementary Table 3). The statistical analyses of IHC and smFISH (RNAscope) density data were carried out in R environment (R version 4.0.2.). We used the ‘ggplot2’ (https://www.rdocumentation.org/packages/ggplot2/versions/3.3.2) and ‘ggthemes’ (https://www.rdocumentation.org/packages/ggthemes/versions/3.5.0) packages for descriptive analysis, ‘nlme’ (https://www.rdocumentation.org/packages/nlme/versions/3.1-150) and ‘multcomp’ (https://www.rdocumentation.org/packages/multcomp/versions/1.4-14) for statistical analysis. To incorporate the unique effect of each donor on which multiple measurements were taken, we applied Linear Mixed Models (LMM) with random effect using the ‘lme’ function of ‘nlme’ package^17–19^. The significance level was set to 1% (α=0.01). Cell types were investigated in separate models. Cell densities were set as dependent variables. Identifiers of individual samples were included as random effect. Layerwise localization (cortical layers 1-6) was set as a 6-level within-subject factor while diagnosis was set as a 2-level between subject factor (schizophrenia or control). The layer*diagnosis interactions indicated the layerwise effect of SCH diagnosis. For direct comparison between specific layers of diagnostic groups, contrast matrices were constructed based on the reported models and tested with the ‘glht’ function of ‘multcomp’ package. In case of significant contrasts, the effects of possible confounders such as age, gender and PMI were tested with correlation analyses. Significance level for normality tests (Shapiro-Wilk test) and correlation tests (Pearson’s – applied in case of normal distribution, Spearman’s – applied when data did not follow normal distribution), was set to the conventional 5% (α=0.05).

No statistically significant interaction between L2 CR+ density and PMI, age or gender was detected. Furthermore, there were no statistically significant morphometric (diameter) differences between the diagnostic groups (Supplementary Table 3). Cortical widths were very similar in schizophrenia and CTR (Supplementary Table 3). While age and gender did not show any significant correlation with the density of L2 PV+ neurons, there was a negative correlation with PMI in controls (r=-0.795, p=0.006) and when groups were combined (r=-0.736, p<0.001).

---

**Extended Data Figure 1.**
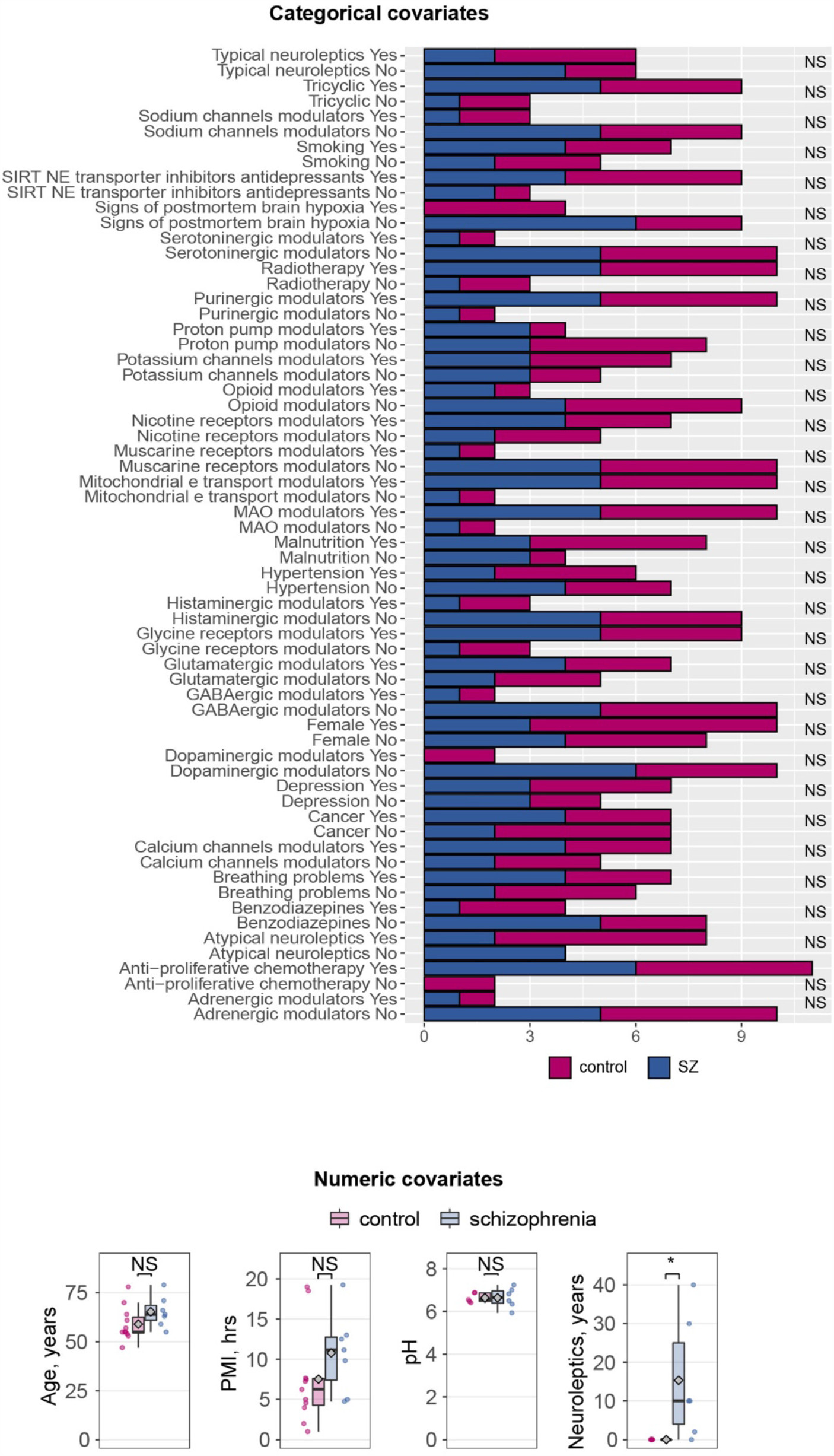
Analysis of potential confounder effect on snRNA-seq dataset from the DLPFC of patients with schizophrenia and control subjects. A: Barplot showing Fisher’s exact test for each covariate with “yes-no” representation, NS – non-significant. B: Comparison of age, PMI (postmortem interval), pH and years of neuroleptic treatment between control and schizophrenia groups. Note an expected difference in years of neuroleptic treatment since schizophrenia, but not control subjects, are usually treated by neuroleptics during the disease course. Normality was tested using Shapiro-Wilk test. Equality of variances was tested using Levene’s test. Significance of differences for age and pH conditions were tested using Student two-tailed T-test; duration of neuroleptic treatment and PMI were tested using Independent 2-group Mann-Whitney U test. * − 0.01 < p ≤ 0.05, NS – non-significant. Horizontal line in the box represents median, diamond represents mean. Lower and upper hinges of the box correspond to the first and third quartiles. Upper and lower whiskers correspond to the smallest and the largest values, and not more than 1.5 x inter-quartile range.

**Extended Data Figure 2.**
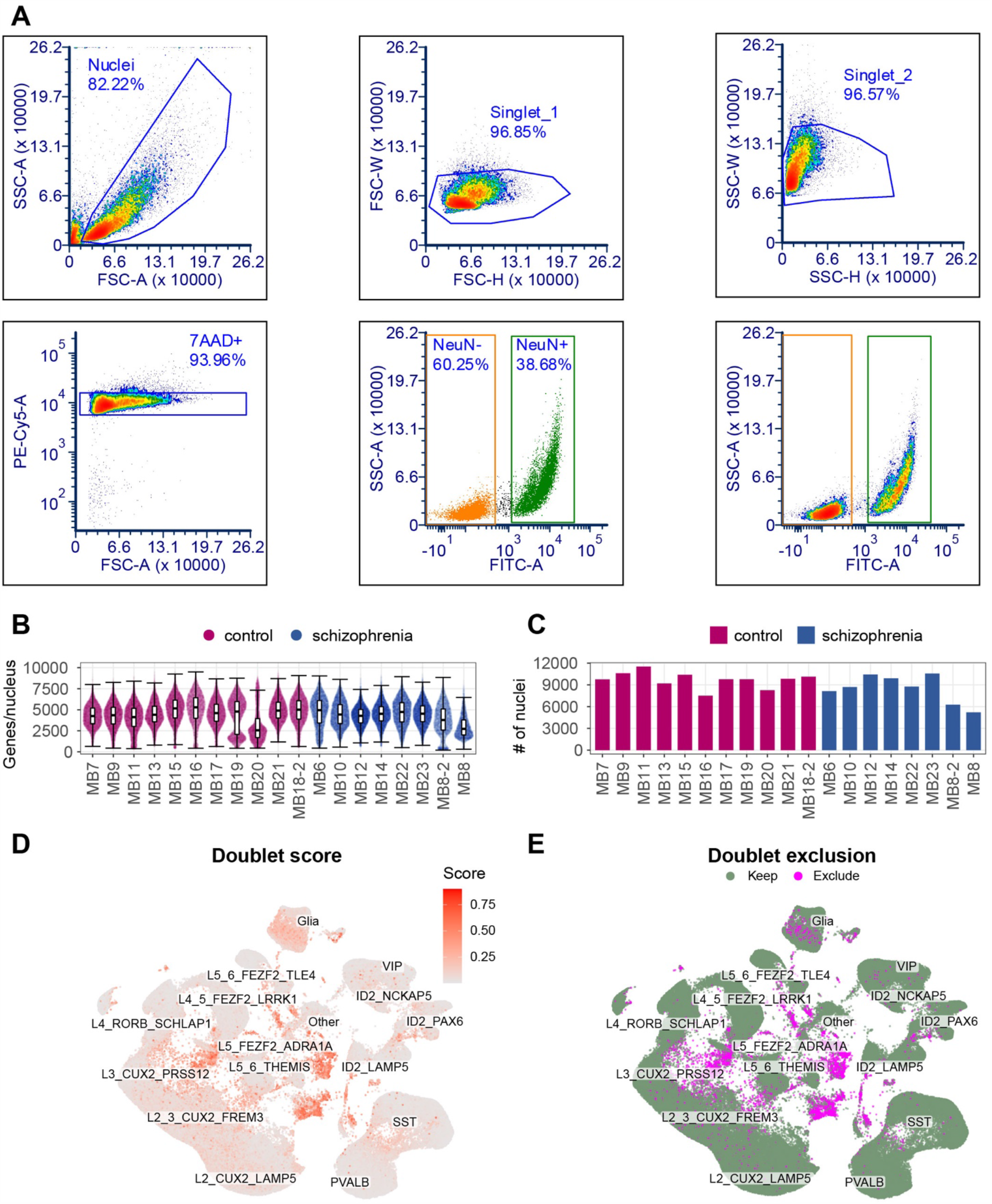
Sorting procedure and initial characterization of snRNA-seq samples. A: Representative FANS plots for flow cytometric isolation of NeuN+ neuronal nuclei from the DLPFC. B: Number of genes per nucleus detected for each sample. Line inside the box represents median, lower and upper hinges of the box correspond to the first and third quartiles. Upper and lower whiskers correspond to the smallest and the largest values, and not more than 1.5 x inter-quartile range. y axis was square-root transformed. C: Number of single nuclei sequenced for each sample. D,E: Identification and exclusion of doublets.

**Extended Data Figure 3.**
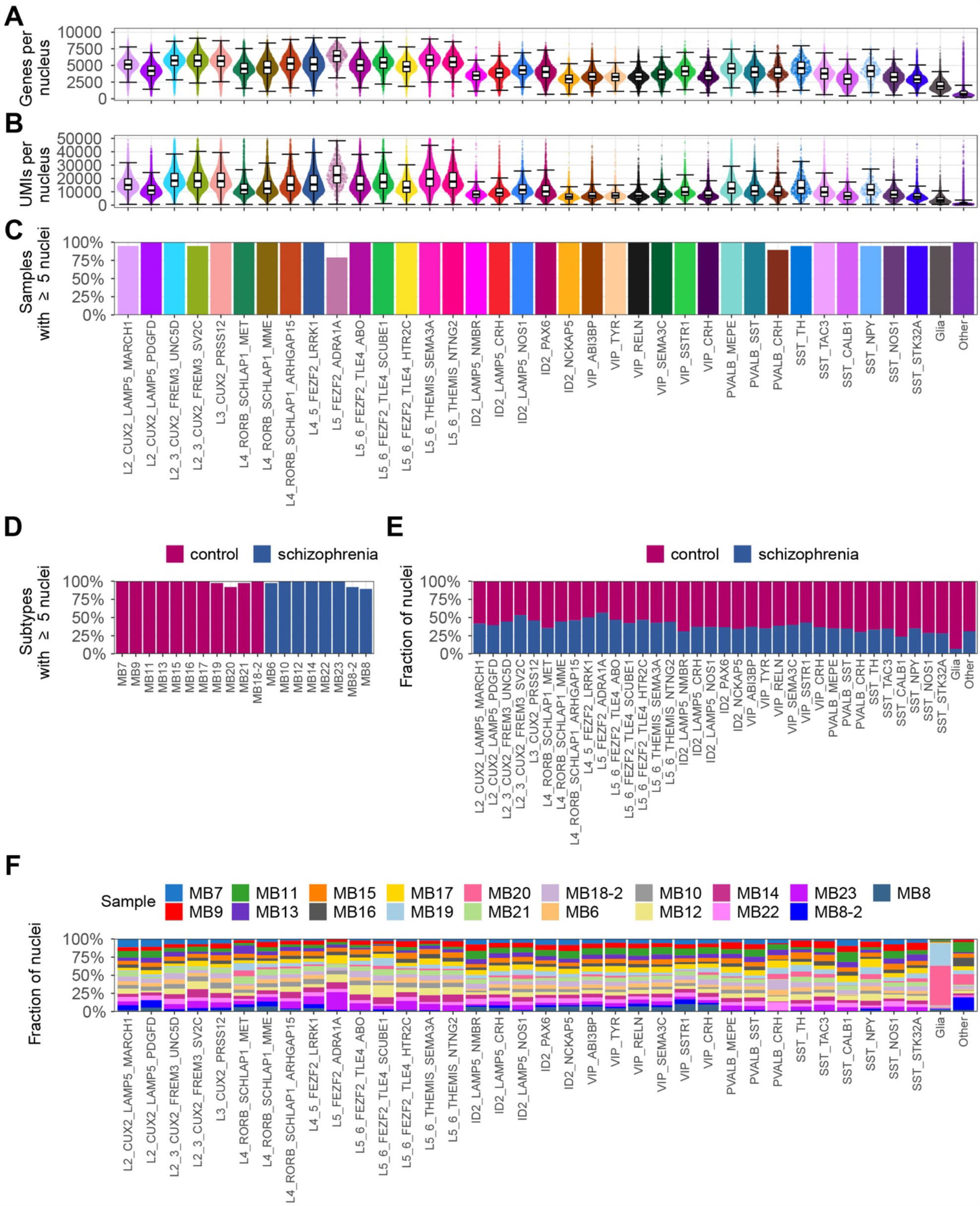
Quality control of snRNA-seq libraries and per-sample analysis of subtype representation. A: Number of genes per nucleus for each subtype. B: Number of UMI per nucleus for each subtype. C: Percentage of samples where a subtype is represented by ≥5 nuclei. D: Percentage of subtypes that were represented by ≥5 nuclei in each sample. E: Fraction of nuclei per subtype shown for conditions. F: Fraction of nuclei per subtype shown for samples. For panels with boxplots: line inside the box represents median, lower and upper hinges of the box correspond to the first and third quartiles. Upper and lower whiskers correspond to the smallest and the largest values, and not more than 1.5 x inter-quartile range.

**Extended Data Figure 4.**
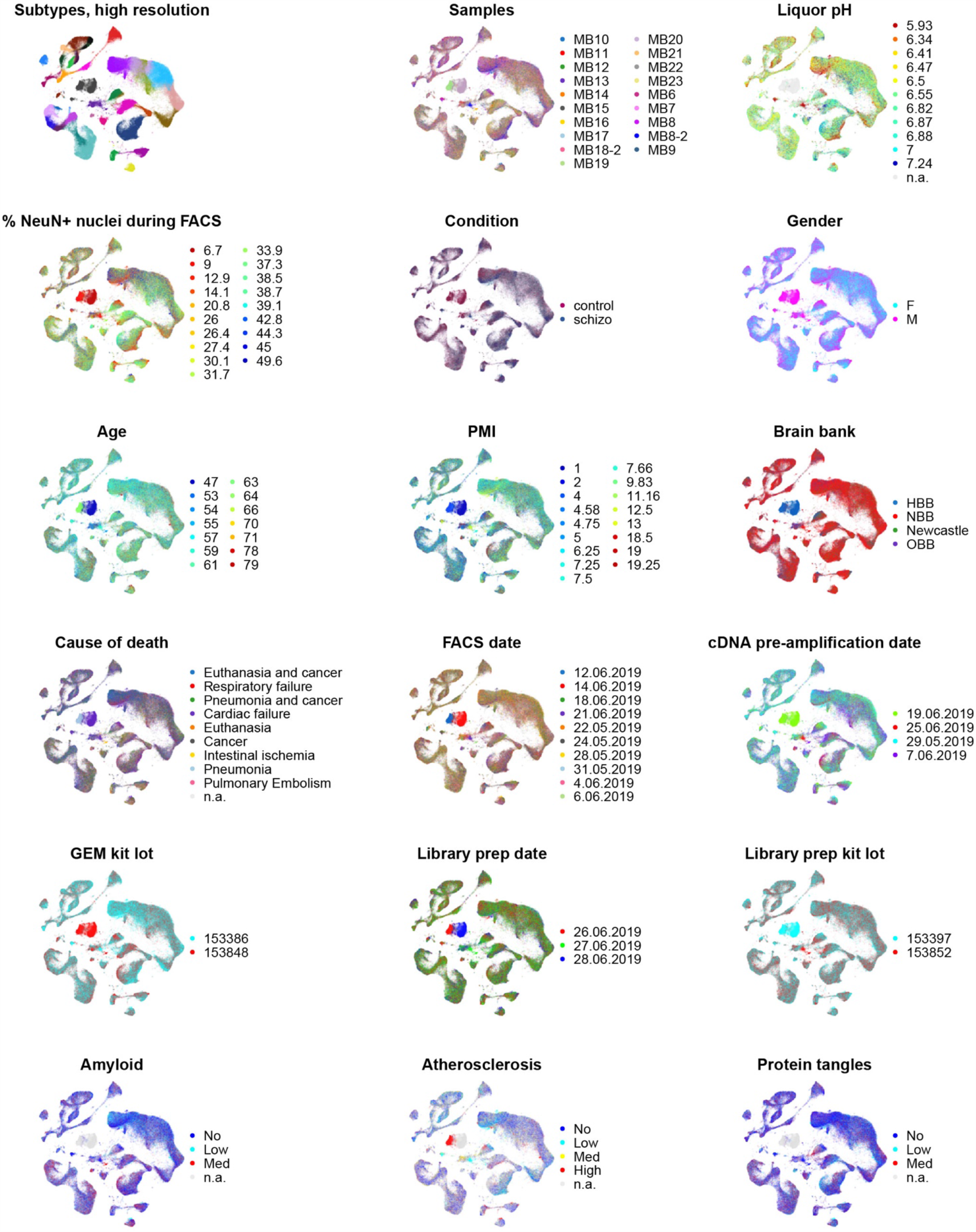
UMAP plots showing the distribution for some of the main covariates and batches of the samples on the common embedding.

**Extended Data Figure 5.**
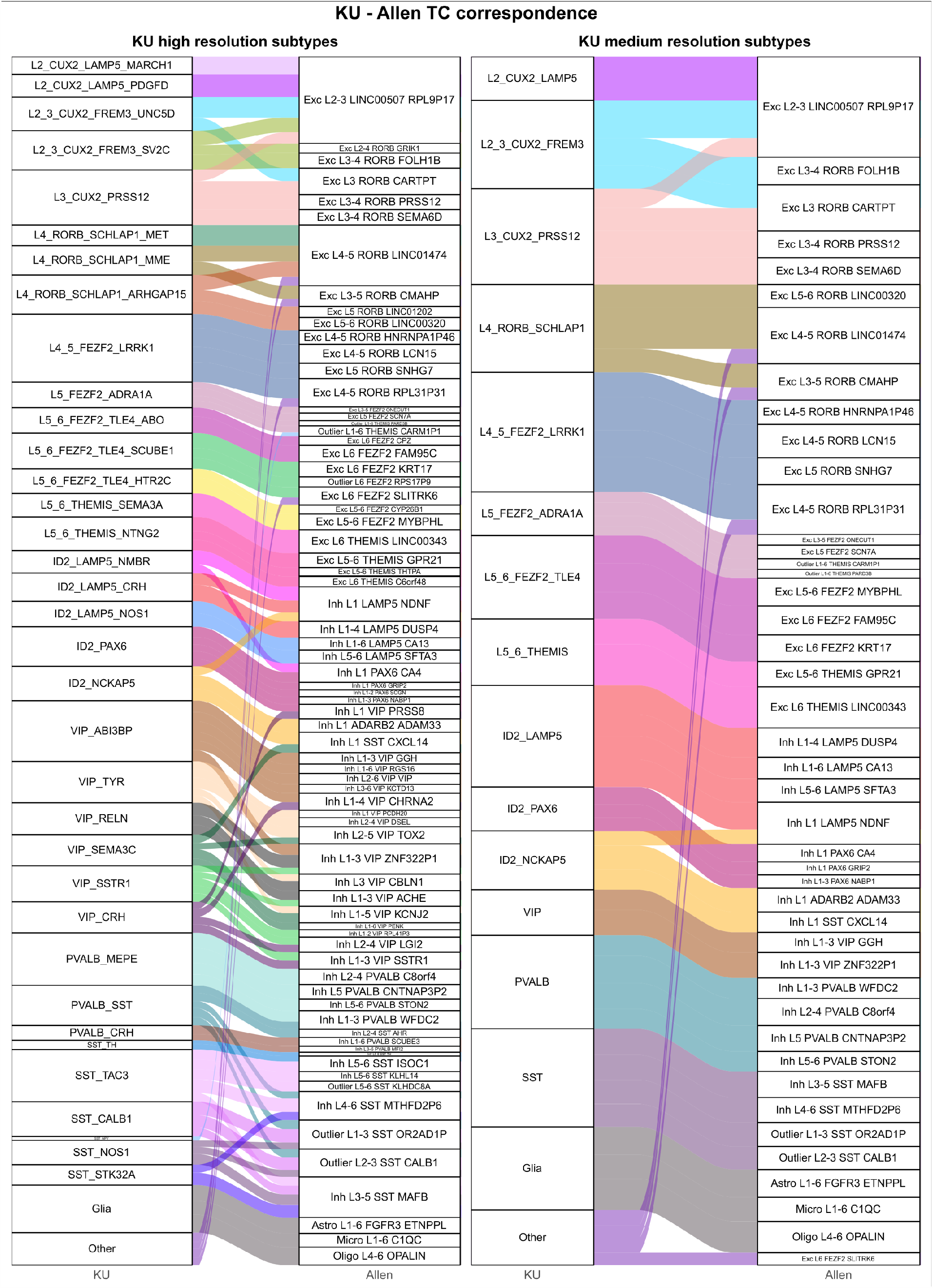
Correspondence of neuronal subtypes from the current dataset with dataset from Allen Brain Institute for normal temporal cortex.

**Extended Data Figure 6.**
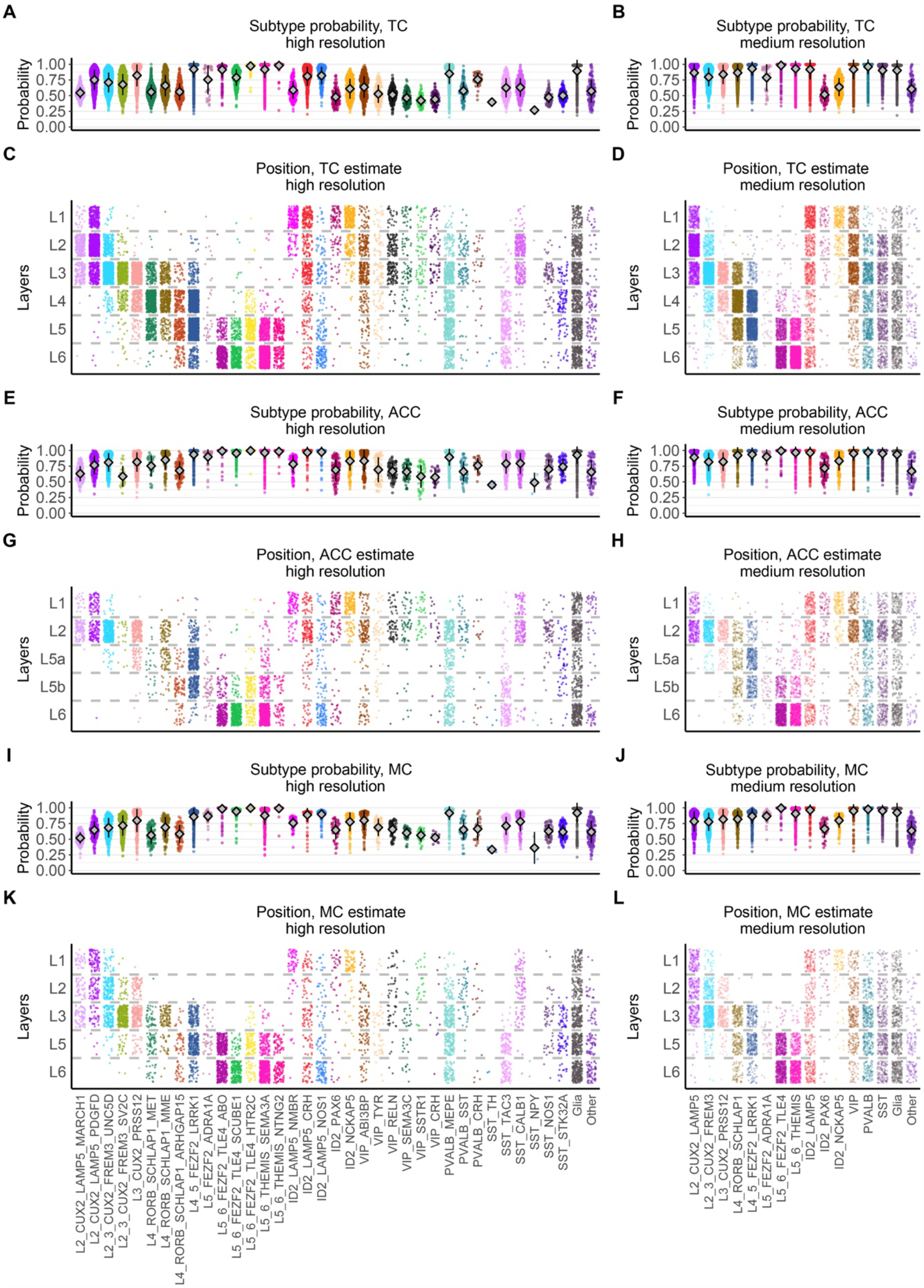
Cross-annotation of layers between the current snRNA-seq dataset of the DLPFC and cortical regions from the normal brain provided by Allen Brain Institute. A-D: Cross annotation of BA9 DLPFC to the layers of the temporal cortex (TC). E-H: Cross annotation of BA9 DLPFC to the layers of the anterior cingulate cortex (ACC). I-L: Cross annotation of BA9 DLPFC to the layers of the motor cortex (ACC). Note that although some differences in subtype composition between BA9 vs TC, ACC, and MC analyzed by Allen Brain Institute were expected, we observed that the layer placement was consistent in most cases.

**Extended Data Figure 7.**
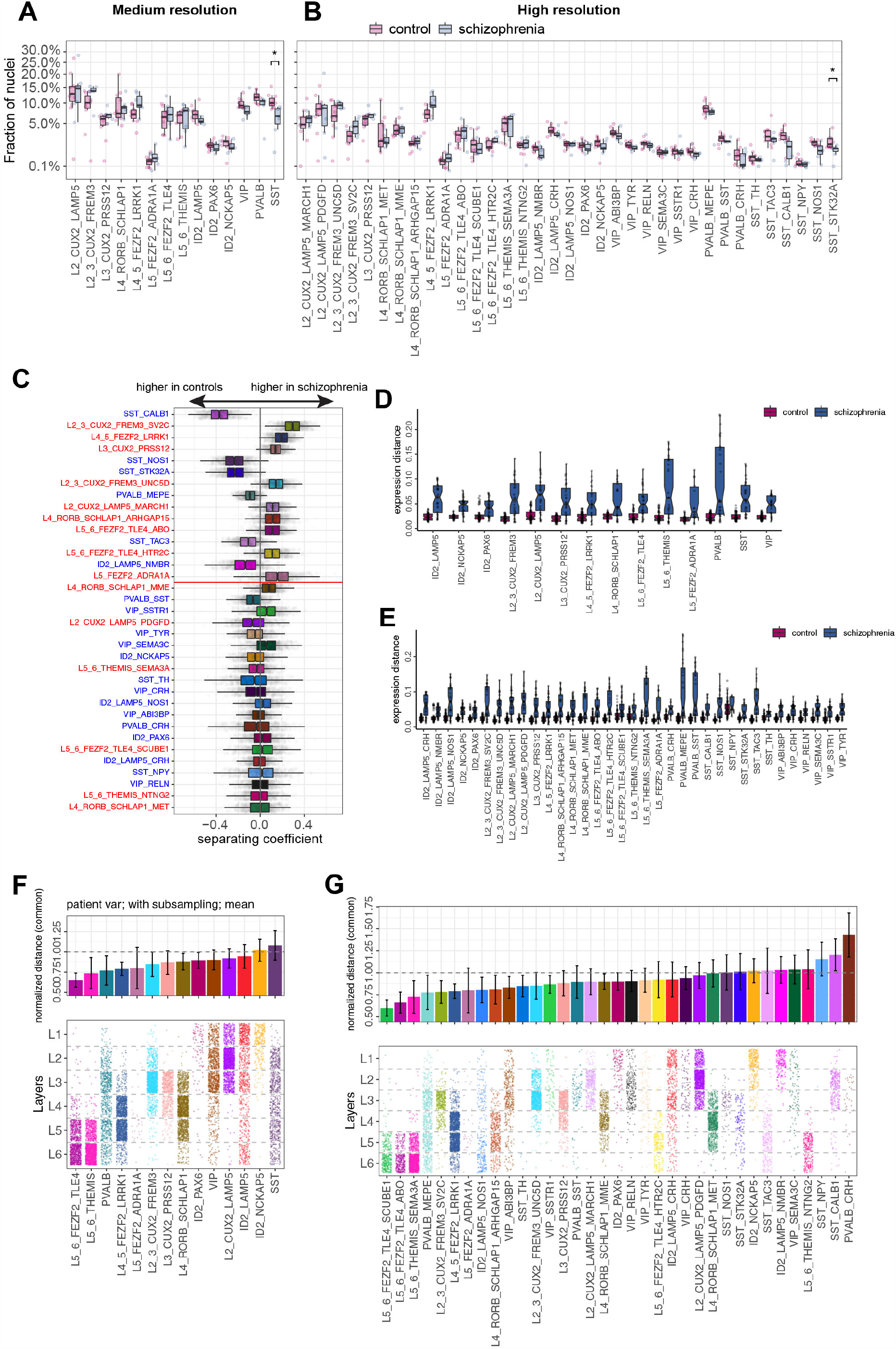
Compositional changes in the DLPFC in schizophrenia identified by snRNA-seq. A,B: Fraction of nuclei represented by each neuronal subtype are shown for the schizophrenia and control groups for medium (A) and high (B) resolutions. Normality was tested using Shapiro-Wilk test, equality of variances was tested using Levene’s test. As part of the data was not normally distributed Independent 2-group Mann-Whitney U test was used to test significance of differences for all group pairs. Multiple comparison correction was done using Benjamini & Hochberg method. p values: SST 0.01, SST_STK32A 0.042. Line inside the box represents median, lower and upper hinges of the box correspond to the first and third quartiles. Upper and lower whiskers correspond to the smallest and the largest values, and not more than 1.5 x inter-quartile range. y axis was square-root transformed. C: As in Fig. 2c of the main text, change in neuronal composition between controls and schizophrenia is shown based on a separating coefficient for all cell types. D,E: Analogous to Fig. 2f, the gene expression distance (based on Pearson correlation) within control and schizophrenia samples is shown for all individual cell types, for medium (D) and high (E) resolutions. F,G: Cell types sorted by the mean expression distance along consensus direction of expression difference between schizophrenia and control samples (see Methods) for medium (F) and high (G) resolution annotations.

**Extended Data Figure 8.**
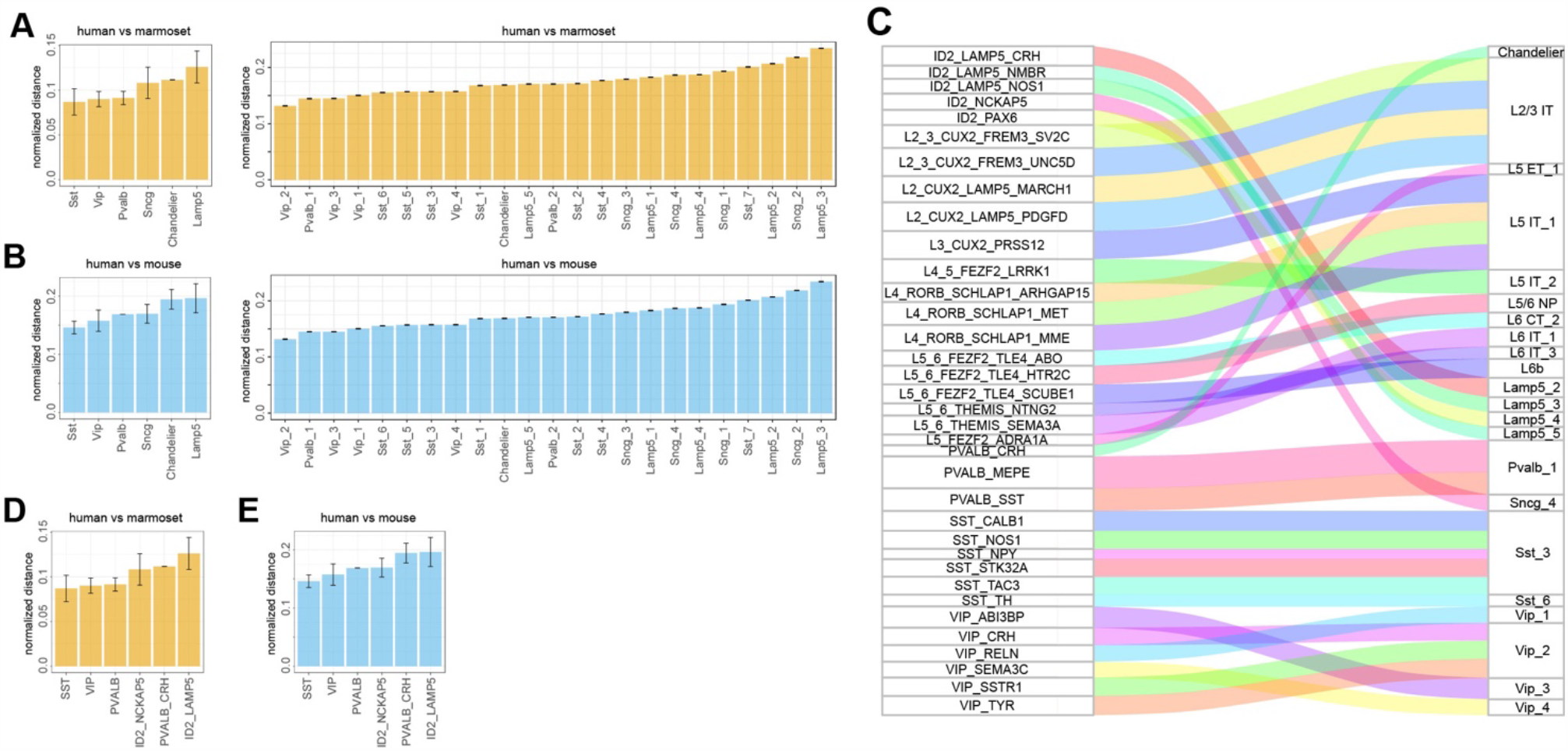
Conservation of gene expression between GABAergic interneuron subtypes of human, marmoset and mouse cortices. A: Gene expression distance (based on Pearson linear correlation) between the orthologous subtypes of GABAergic interneurons in human and marmoset motor cortex (M1 region). Analysis based on medium (left) and high (right) resolutions is shown. Data is taken from BICCN dataset from Bakken et al. 2020, bioRxiv 2020.03.31.016972. B: Same as A, comparing human and mouse motor cortex (M1 region). C: Correspondence of the neuronal subtype annotation between the current dataset and Bakken et al. 2020. D: Gene expression distance between correspondent subtypes of GABAergic interneurons from human and marmoset primary motor cortex with annotation labels from the current dataset. E: Same as D for human and mouse.

**Extended Data Figure 9.**
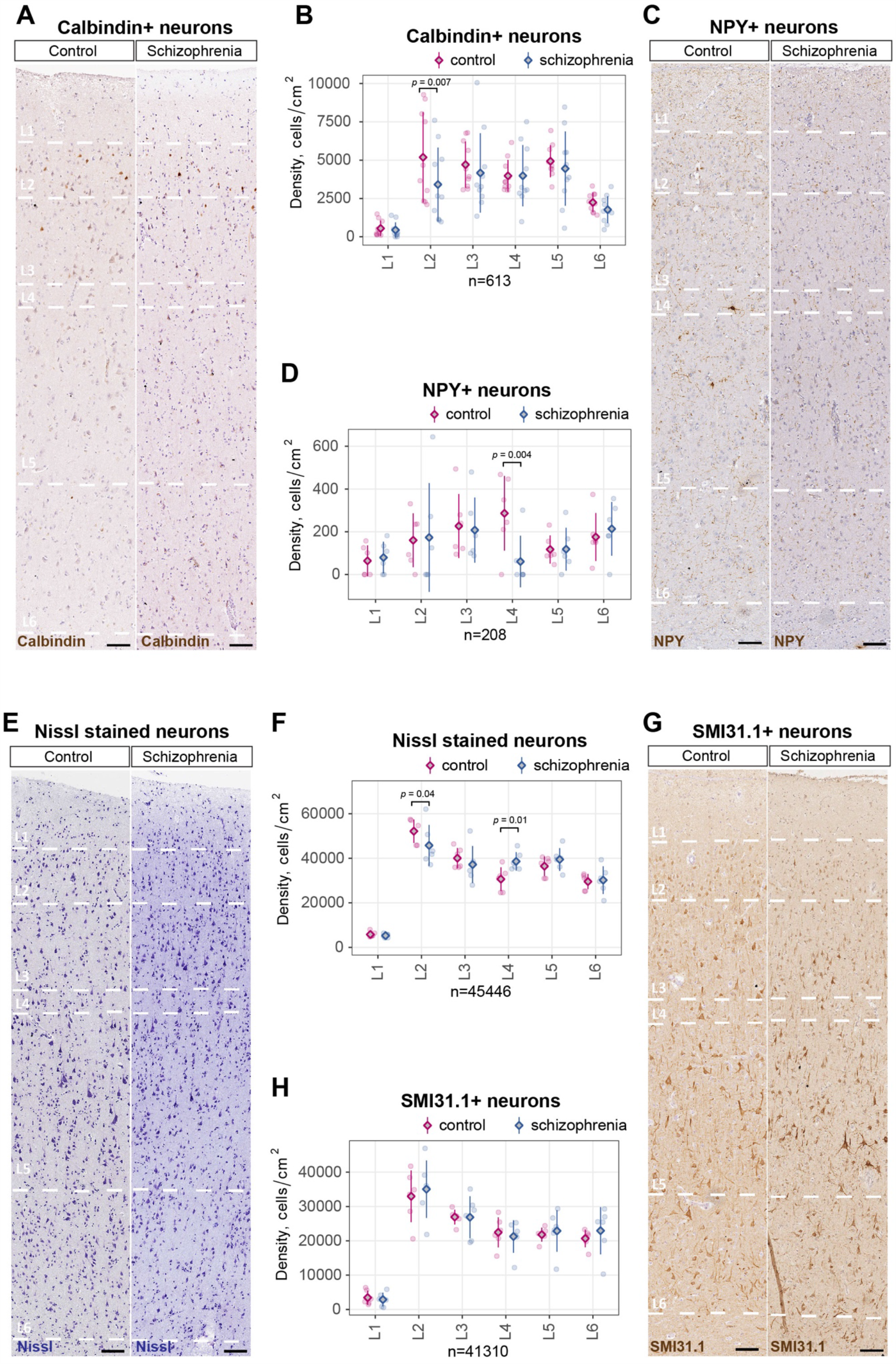
Changes in density and distribution of neuronal subtypes in the DLPFC in schizophrenia analyzed by immunohistochemistry. A,B: Representative images and layer-wise quantification of GABAergic interneurons that express calbindin in the DLPFC in patients with schizophrenia and control subjects. C,D: Representative images and layer-wise quantification of GABAergic interneurons that express neuropeptide Y (NPY) neurons in the DLPFC in patients with schizophrenia and control subjects. E,F: Representative images and layer-wise quantification of Nissl labeled neurons in the DLPFC in patients with schizophrenia and control subjects. G,H: Representative images and layer-wise quantification of principal neurons labeled by SMI31.1 in the DLPFC in patients with schizophrenia and control subjects. Diamonds on the plots represent mean, error bars +-SD.

**Extended Data Figure 10.**
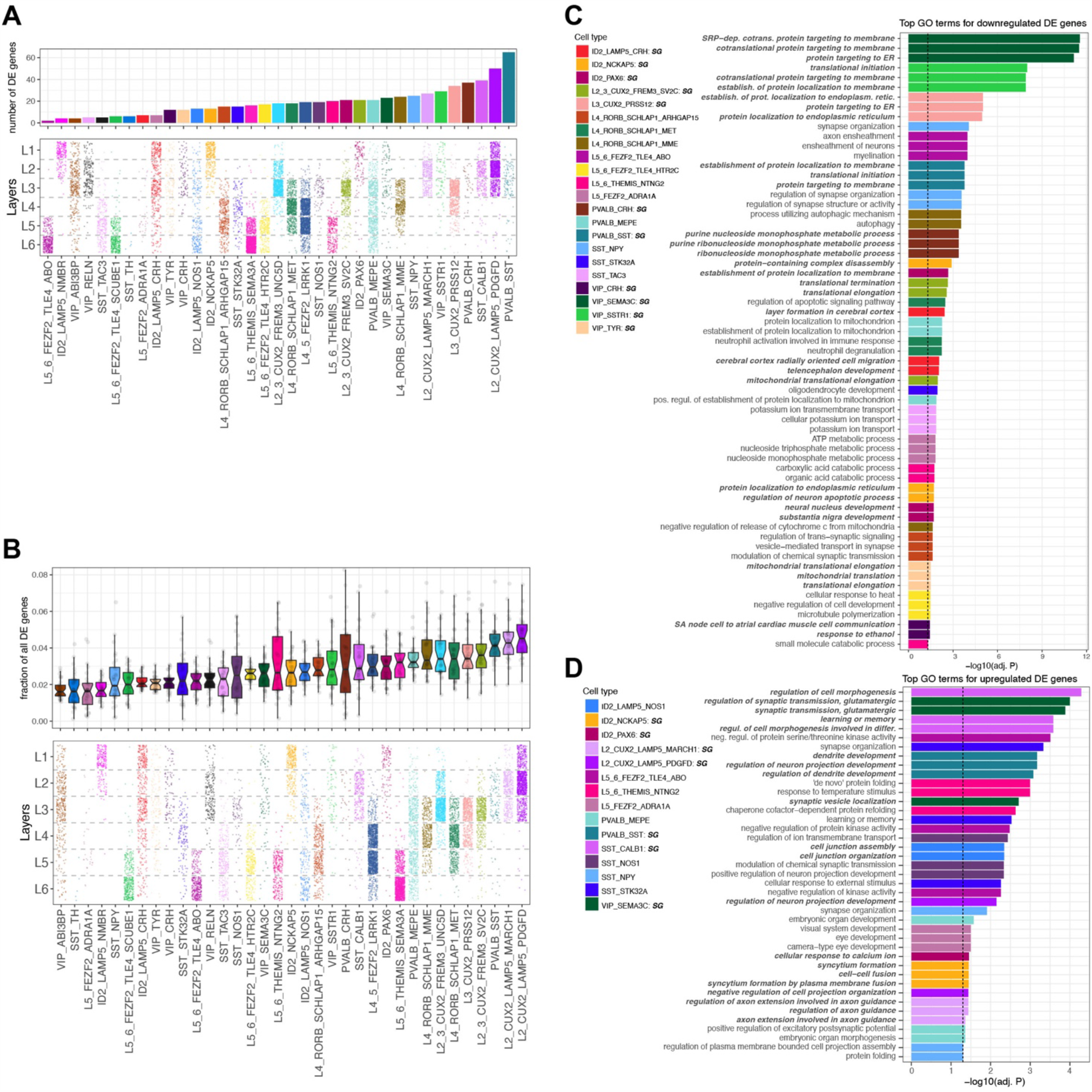
Schizophrenia-associated changes in the differentially expressed (DE) genes and enriched GO terms for each subtype. A: Number of DE genes between schizophrenia and control groups shown for each neuronal subtype. For this test, a fixed number of cells was sampled from each cell type. The bottom plot shows predicted cortical layer positions. B: Fraction of all DE genes represented by each subtype, evaluated using bootstrap resampling of available samples within the schizophrenia and control samples, as well as sampling a fixed number of cells from each cell type. C,D: Full list of GO terms significantly enriched in the set of top down- and upregulated genes in the neuronal subtypes from schizophrenia DLPFC, compared to controls.

**Extended Data Figure 11.**
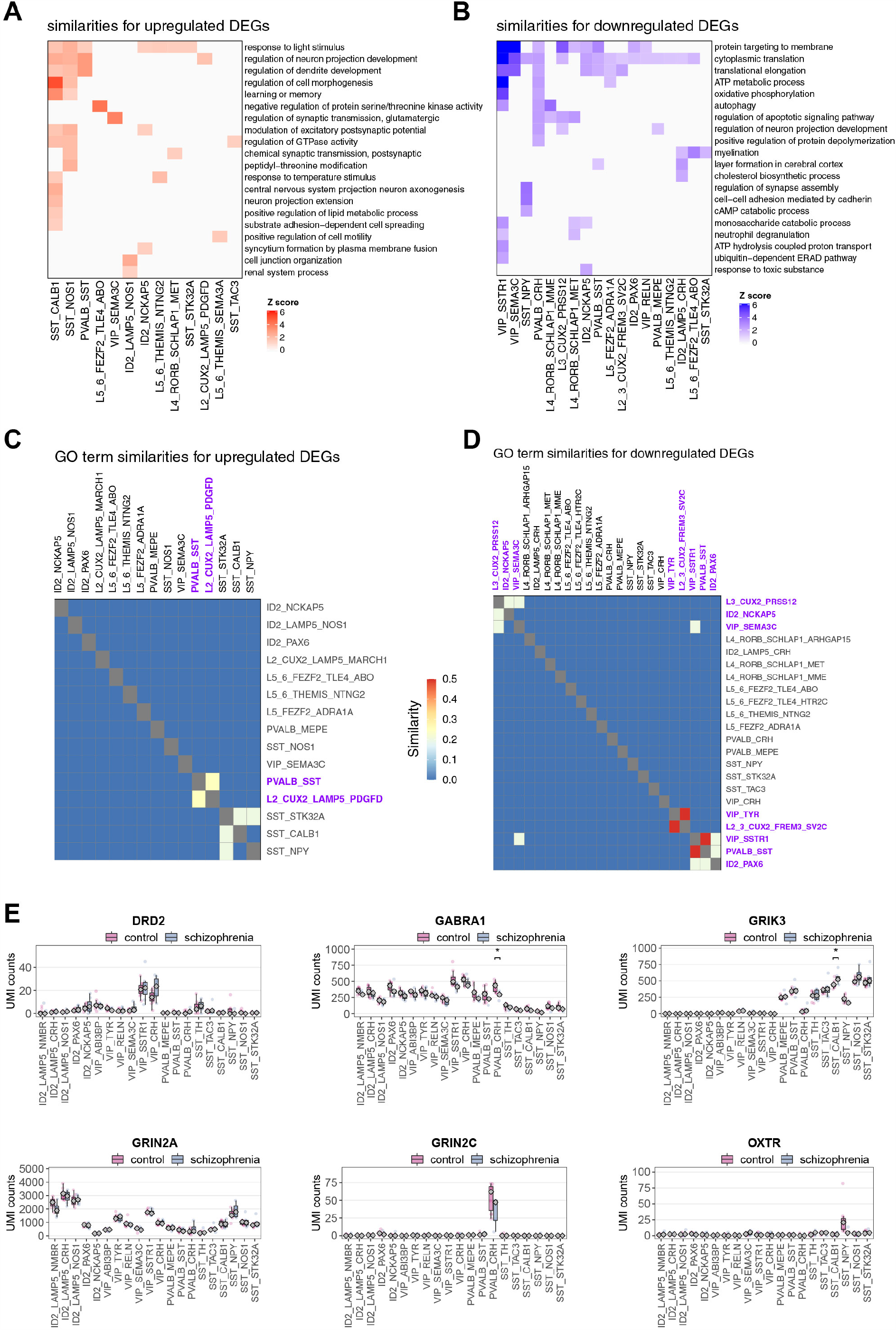
Correlation of gene expression changes across neuronal subtypes in the DLPFC in schizophrenia. A,B: Heatmaps showing clusters of GO terms significantly (BH adjusted P value of 0.05 or lower) enriched in the set of upregulated (A) and downregulated (B) genes between schizophrenia and controls. Each row corresponds to a set of related GO terms (clustered based on the gene content), with each column representing a separate cell type. C,D: Heatmaps showing clustering of neuronal subtypes based on similarities of GO terms enriched in genes upregulated (C) or downregulated (D) in schizophrenia relative to controls. E: Schizophrenia-related change of genes that code receptors that were previously proposed as targets for schizophrenia drugs in inhibitory neuronal subtypes. p values were estimated using differential gene expression Wald test in DESeq2. Multiple comparison correction was done using Benjamini & Hochberg method. p values: GABRA1 0.0346733, GRIK3 0.03965785. Expression levels are based on normalized pseudo-bulk mRNA UMI counts. Diamond represents median.

**Extended Data Figure 12.**
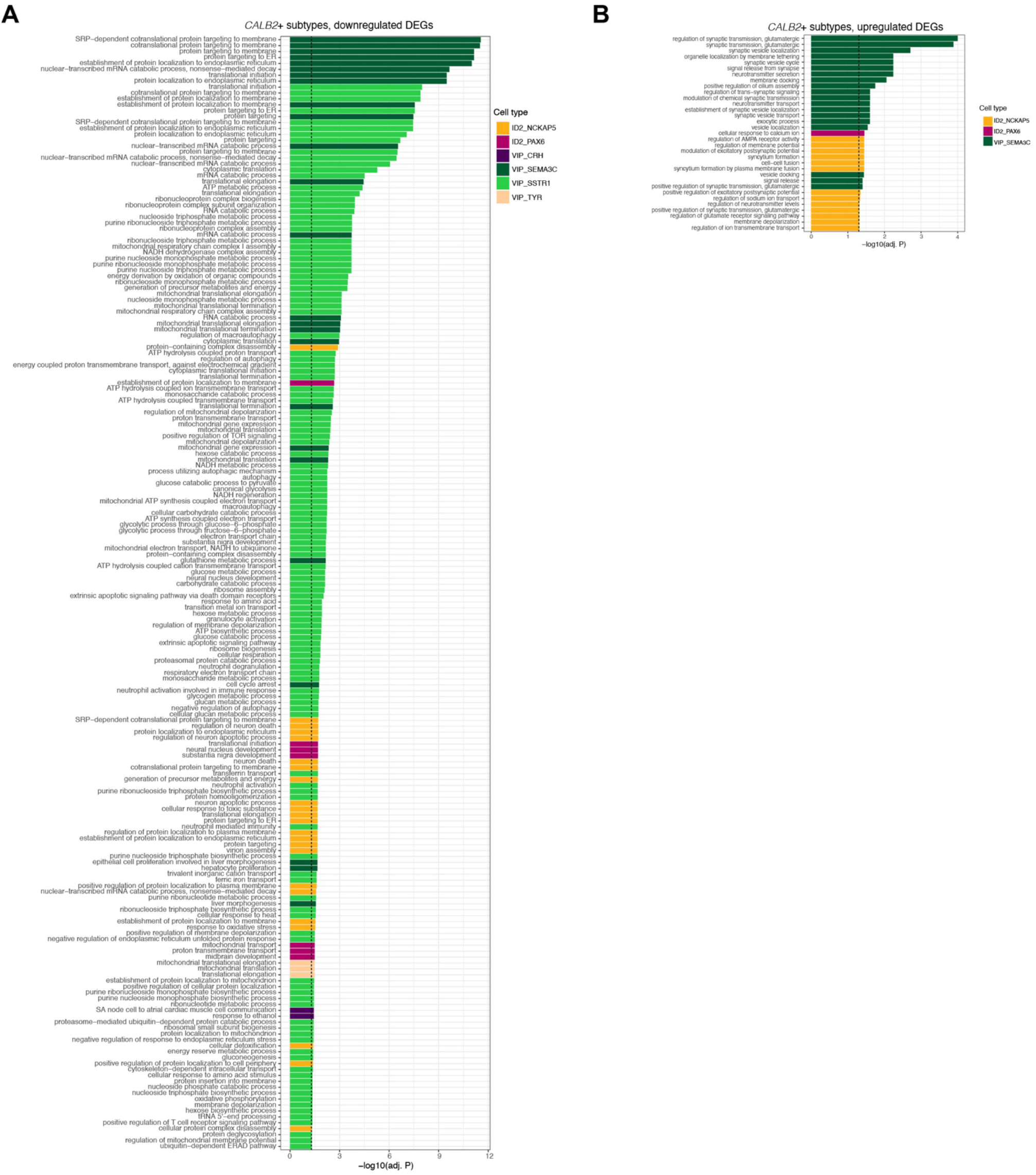
GO terms enriched in schizophrenia-associated changes for neuronal subtypes of VIP and ID2 families that express CALB2. A,B: Full list of significantly enriched GO terms in genes that were downregulated (A) or upregulated (B) in the CALB2+ neuronal subtypes of schizophrenia DLPFC relative to controls.

**Extended Data Figure 13.**
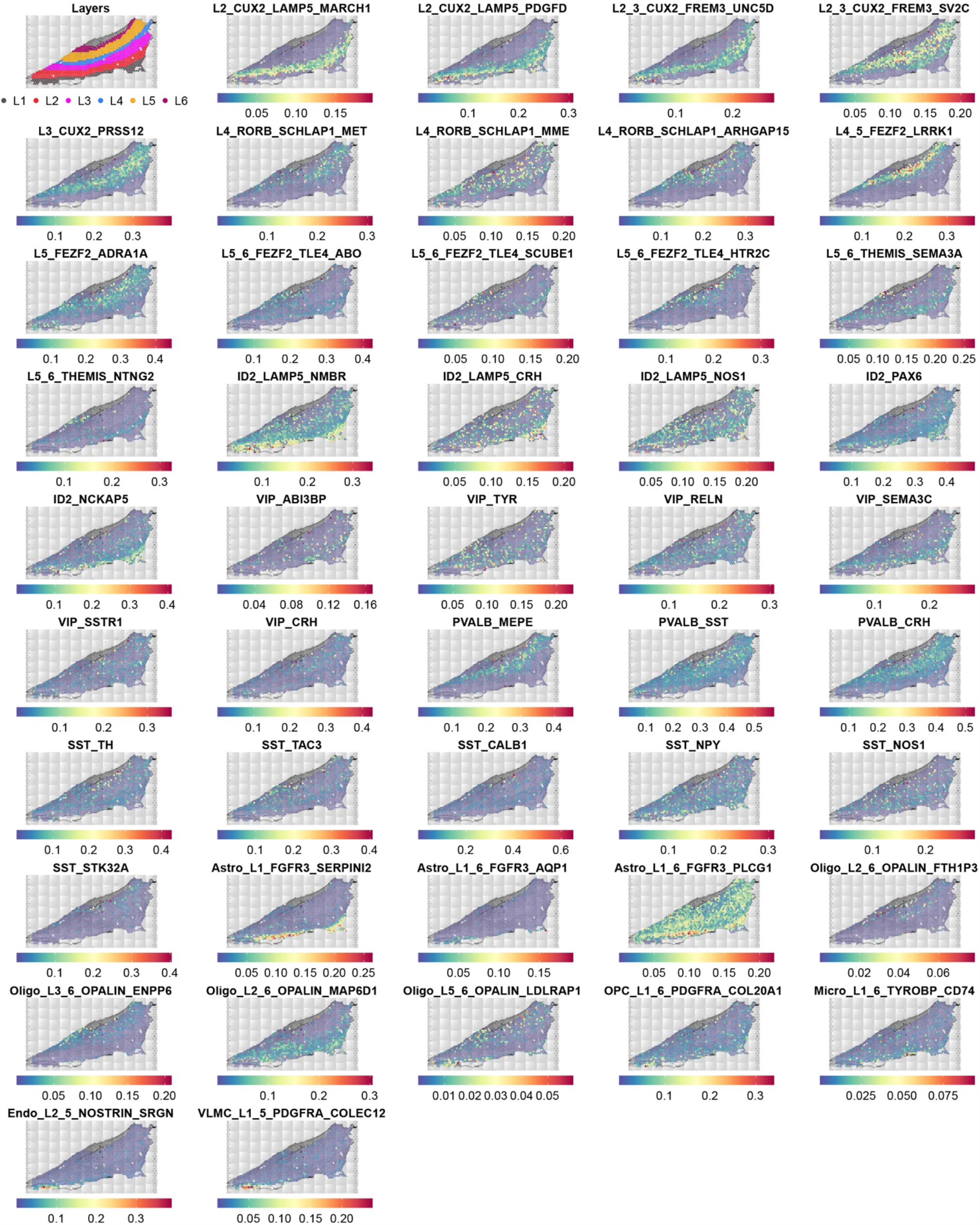
Distribution of neural cell subtypes in the DLPFC by spatial transcriptomics using markers previously identified in the current snRNA-seq dataset. Cortical layers represent manual annotation of the tissue by experienced neuroanatomist. Each spot on Visium slide is colored by the fraction of predicted neural cell subtypes in the overlaying tissue slice.

**Extended Data Figure 14.**
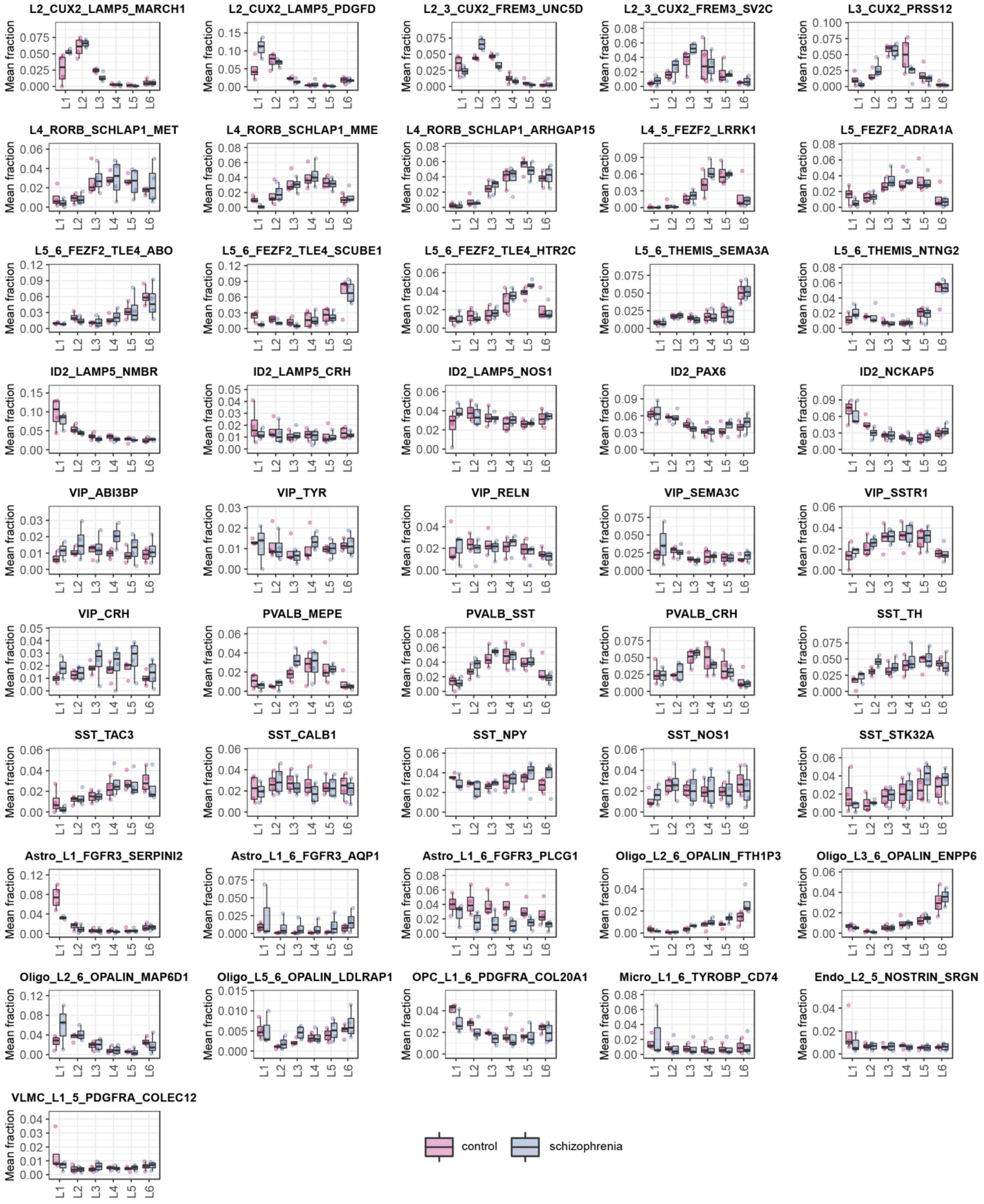
Changes in neural cell composition in the DLPFC in schizophrenia identified by spatial transcriptomics across separate cortical layers. Fraction of separate neural subtypes was averaged across separate cortical layers for each sample. Normality was tested using Shapiro-Wilk test, equality of variances was tested using Levene’s test. As part of the data was not normally distributed Independent 2-group Mann-Whitney U test was used to test significance of differences for all group pairs. Multiple comparison correction was done using Benjamini & Hochberg method. Line inside the box represents median, lower and upper hinges of the box correspond to the first and third quartiles. Upper and lower whiskers correspond to the smallest and the largest values, and not more than 1.5 x inter-quartile range.

**Extended Data Figure 15.**
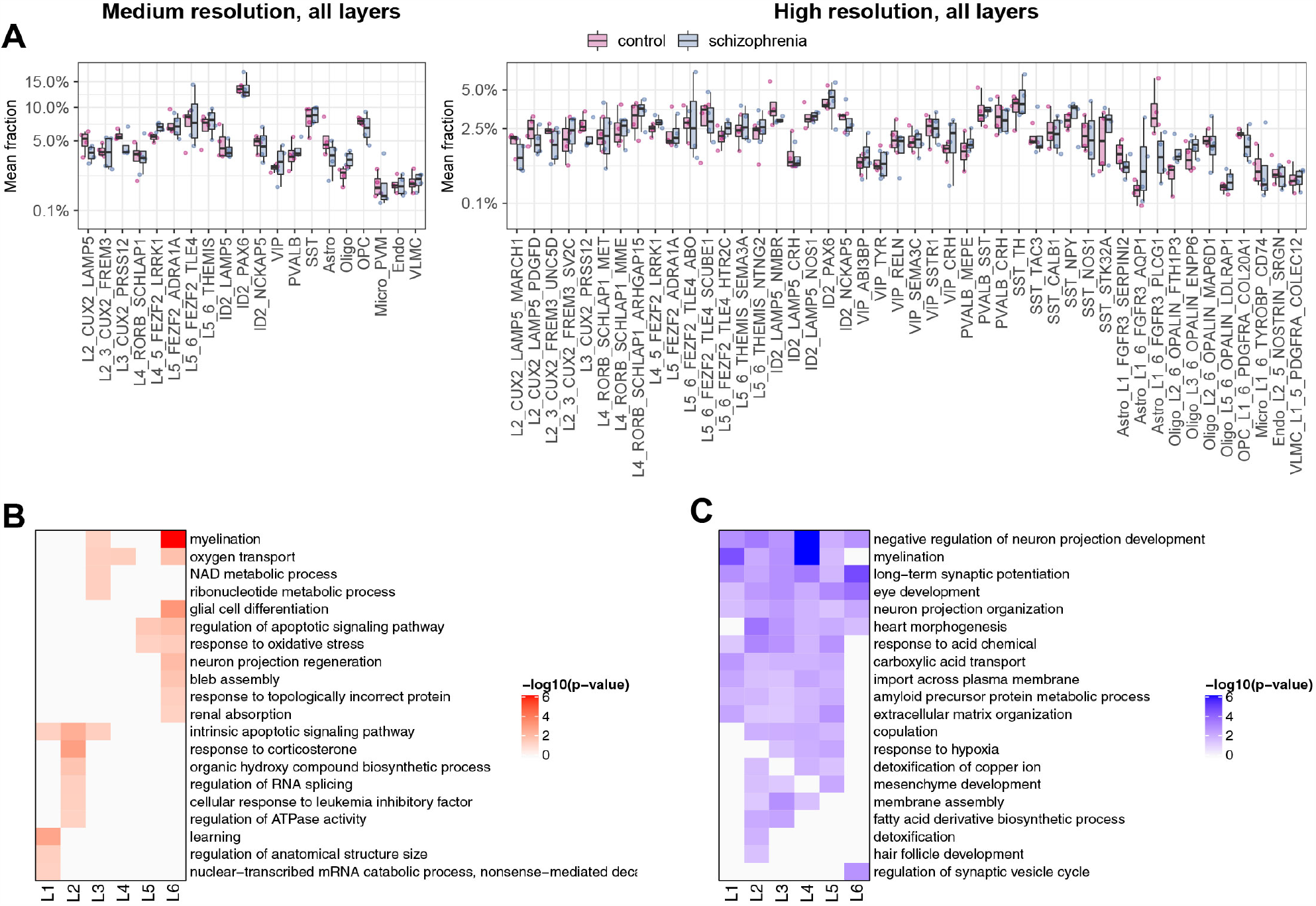
Analysis of schizophrenia-associated changes in neural cell fractions and GO terms by spatial transcriptomics. A,B: Fraction of neural cells (in %) for each subtype between control and schizophrenia samples at medium and high resolution across all cortical layers. Fraction of separate neural subtypes was averaged across whole sample. Normality was tested using Shapiro-Wilk test, equality of variances was tested using Levene’s test. As part of the data was not normally distributed Independent 2-group Mann-Whitney U test was used to test significance of differences for all group pairs. Multiple comparison correction was done using Benjamini & Hochberg method. Line inside the box represents median, lower and upper hinges of the box correspond to the first and third quartiles. Upper and lower whiskers correspond to the smallest and the largest values, and not more than 1.5 x inter-quartile range. y axis was square-root transformed. C,D: Spatial GO term groups clustered based on the similarity of the up-regulated (B) and down-regulated (C) differentially expressed genes. p values represent Benjamini-Hochberg corrected values of 0.05 or lower.

## REFERENCES

1. Millan, M. J. et al. Altering the course of schizophrenia: progress and perspectives. Nat. Rev. Drug Discov. 15, 485–515 (2016).

2. Owen, M. J., Sawa, A. & Mortensen, P. B. Schizophrenia. Lancet 388, 86–97 (2016).

3. Volk, D. W., Edelson, J. R. & Lewis, D. A. Altered expression of developmental regulators of parvalbumin and somatostatin neurons in the prefrontal cortex in schizophrenia. Schizophr Res 177, 3–9 (2016).

4. Borkowska, M., Millar, J. K. & Price, D. J. Altered Disrupted-in-Schizophrenia-1 Function Affects the Development of Cortical Parvalbumin Interneurons by an Indirect Mechanism. PLoS One 11, e0156082 (2016).

5. Glausier, J. R., Fish, K. N. & Lewis, D. A. Altered parvalbumin basket cell inputs in the dorsolateral prefrontal cortex of schizophrenia subjects. Mol Psychiatry 19, 30–36 (2014).

6. Reynolds, G. P. & Beasley, C. L. GABAergic neuronal subtypes in the human frontal cortex - Development and deficits in schizophrenia. J. Chem. Neuroanat. 22, 95–100 (2001).

7. Fromer, M. et al. Gene expression elucidates functional impact of polygenic risk for schizophrenia. Nat Neurosci 19, 1442–1453 (2016).

8. Bowen, E. F. W., Burgess, J. L., Granger, R., Kleinman, J. E. & Rhodes, C. H. DLPFC transcriptome defines two molecular subtypes of schizophrenia. Transl. Psychiatry 9, 1–10 (2019).

9. Fillman, S. G. et al. Increased inflammatory markers identified in the dorsolateral prefrontal cortex of individuals with schizophrenia. Mol Psychiatry 18, 206–214 (2013).

10. Gandal, M. J. et al. Shared molecular neuropathology across major psychiatric disorders parallels polygenic overlap. Science (80-.). 359, 693–697 (2018).

11. Pierri, J. N., Volk, C. L. E., Auh, S., Sampson, A. & Lewis, D. A. Decreased somal size of deep layer 3 pyramidal neurons in the prefrontal cortex of subjects with schizophrenia. Arch. Gen. Psychiatry 58, 466–473 (2001).

12. Holmes, A. J. et al. Prefrontal functioning during context processing in schizophrenia and major depression: An event-related fMRI study. Schizophr. Res. 76, 199–206 (2005).

13. MacDonald, A. W. et al. Specificity of prefrontal dysfunction and context processing deficits to schizophrenia in never-medicated patients with first-episode psychosis. Am. J. Psychiatry 162, 475–484 (2005).

14. Barkas, N. et al. Joint analysis of heterogeneous single-cell RNA-seq dataset collections. Nat. Methods 16, 695–698 (2019).

15. Pfisterer, U. et al. Identification of epilepsy-associated neuronal subtypes and gene expression underlying epileptogenesis. Nat. Commun. 11, 1–19 (2020).

16. Hodge, R. D. et al. Conserved cell types with divergent features in human versus mouse cortex. Nature 573, 61–68 (2019).

17. Allen Institute Cell Types Database: RNA-Seq Data; Multiple Cortical Areas - SMART-SEQ (2019).

18. Baryawno, N. et al. Impact of metastatic prostate cancer on human bone marrow. bioRxiv 2020.03.19.998658 (2020) doi:10.1101/2020.03.19.998658.

19. Defelipe, J. The evolution of the brain, the human nature of cortical circuits, and intellectual creativity. Front. Neuroanat. 5, 29 (2011).

20. Mohan, H. et al. Dendritic and axonal architecture of individual pyramidal neurons across layers of adult human neocortex. Cereb. Cortex 25, 4839–4853 (2015).

21. Berg, J. et al. Human cortical expansion involves diversification and specialization of supragranular intratelencephalic-projecting neurons. bioRxiv 2020.03.31.018820 (2020) doi:10.1101/2020.03.31.018820.

22. Krienen, F. M. et al. Innovations present in the primate interneuron repertoire. Nature 1–8 (2020) doi:10.1038/s41586-020-2781-z.

23. Bakken, T. E. et al. Evolution of cellular diversity in primary motor cortex of human, marmoset monkey, and mouse. bioRxiv 2020.03.31.016972 (2020) doi:10.1101/2020.03.31.016972.

24. Hladnik, A., Džaja, D., Darmopil, S., Jovanov-Milošević, N. & Petanjek, Z. Spatio-temporal extension in site of origin for cortical calretinin neurons in primates. Front. Neuroanat. 8, (2014).

25. Chung, D. W., Fish, K. N. & Lewis, D. A. Pathological Basis for Deficient Excitatory Drive to Cortical Parvalbumin Interneurons in Schizophrenia. Am. J. Psychiatry appiajp201616010025 (2016) doi:10.1176/appi.ajp.2016.16010025.

26. Vasistha, N. A. et al. Maternal inflammation has a profound effect on cortical interneuron development in a stage and subtype-specific manner. Mol. Psychiatry 25, 2313–2329 (2020).

27. Hashimoto, T. et al. Alterations in GABA-related transcriptome in the dorsolateral prefrontal cortex of subjects with schizophrenia. Mol. Psychiatry 13, 147–161 (2008).

28. Montag, C. et al. Association between Oxytocin Receptor Gene Polymorphisms and Self-Rated ‘Empathic Concern’ in Schizophrenia. PLoS One 7, e51882 (2012).

29. Sinkus, M. L. et al. The human CHRNA7 and CHRFAM7A genes: A review of the genetics, regulation, and function. Neuropharmacology vol. 96 274–288 (2015).

30. Sousa, A. M. M., Meyer, K. A., Santpere, G., Gulden, F. O. & Sestan, N. Evolution of the Human Nervous System Function, Structure, and Development. Cell vol. 170 226–247 (2017).

31. Dehay, C. & Kennedy, H. Evolution of the human brain. Science (80-.). 369, (2020).

32. Marín-Padilla, M. The mammalian neocortex new pyramidal neuron: a new conception. Front. Neuroanat. 7, 51 (2014).

33. Wang, D. et al. Comprehensive functional genomic resource and integrative model for the human brain. Science (80-.). 362, eaat8464 (2018).

34. Li, M. et al. Integrative functional genomic analysis of human brain development and neuropsychiatric risks. Science 362, eaat7615 (2018).

35. Skene, N. G. et al. Genetic identification of brain cell types underlying schizophrenia. Nat. Genet. 50, 825–833 (2018).

36. Petanjek, Z. et al. The Protracted Maturation of Associative Layer IIIC Pyramidal Neurons in the Human Prefrontal Cortex During Childhood: A Major Role in Cognitive Development and Selective Alteration in Autism. Front. Psychiatry 10, 122 (2019).

37. Zecevic, N. & Rakic, P. Development of layer I neurons in the primate cerebral cortex. J. Neurosci. 21, 5607–5619 (2001).

38. Silbereis, J. C., Pochareddy, S., Zhu, Y., Li, M. F. & Sestan, N. The Cellular and Molecular Landscapes of the Developing Human Central Nervous System. Neuron 89, 248–268 (2016).

39. Petanjek, Z. et al. Extraordinary neoteny of synaptic spines in the human prefrontal cortex. Proc. Natl. Acad. Sci. U. S. A. 108, 13281–13286 (2011).

40. Sanz-Morello, B. et al. Complex IV subunit isoform COX6A2 protects fast-spiking interneurons from oxidative stress and supports their function. EMBO J. 39, e105759 (2020).

41. Velmeshev, D. et al. Single-cell genomics identifies cell type–specific molecular changes in autism. Science (80-.). 364, 685–689 (2019).

## Supplementary references

1. Bakken, T. E. et al. Evolution of cellular diversity in primary motor cortex of human, marmoset monkey, and mouse. bioRxiv 2020.03.31.016972 (2020) doi:10.1101/2020.03.31.016972.

2. Baryawno, N. et al. Impact of metastatic prostate cancer on human bone marrow. bioRxiv 2020.03.19.998658 (2020) doi:10.1101/2020.03.19.998658.

3. Love, M. I., Huber, W. & Anders, S. Moderated estimation of fold change and dispersion for RNA-seq data with DESeq2. Genome Biol. 15, 550 (2014).

4. Barkas, N. et al. Joint analysis of heterogeneous single-cell RNA-seq dataset collections. Nat. Methods 16, 695–698 (2019).

5. Crowell, H. L. et al. On the discovery of population-specific state transitions from multisample multi-condition single-cell RNA sequencing data. bioRxiv 713412 (2019) doi:10.1101/713412.

6. Yu, G., Wang, L.-G., Han, Y. & He, Q.-Y. clusterProfiler: an R Package for Comparing Biological Themes Among Gene Clusters. Omi. A J. Integr. Biol. 16, 284–287 (2012).

7. Allen Institute Cell Types Database: RNA-Seq Data; Multiple Cortical Areas - SMART-SEQ (2019).

8. Hodge, R. D. et al. Conserved cell types with divergent features in human versus mouse cortex. Nature 573, 61–68 (2019).

9. Pertea, G. & Pertea, M. GFF Utilities: GffRead and GffCompare. F1000Research 9, 304 (2020).

10. Schindelin, J. et al. Fiji: an open-source platform for biological-image analysis. Nat Methods 676–682 (2012).

11. (BICCN), B. I. C. C. N. et al. Title: A multimodal cell census and atlas of the mammalian primary motor cortex Authors: BRAIN Initiative Cell Census Network (BICCN). bioRxiv 2020.10.19.343129 (2020) doi:10.1101/2020.10.19.343129.

12. Stuart, T. et al. Comprehensive Integration of Single-Cell Data. Cell 177, 1888-1902.e21 (2019).

13. Adorjan, I. et al. Calretinin interneuron density in the caudate nucleus is lower in autism spectrum disorder. Brain 140, 2028–2040 (2017).

14. Petrides, M. Dorsolateral prefrontal cortex: Comparative cytoarchitectonic analysis”in the human and the macaque brain and corticocortical connection patterns. Eur. J. Neurosci. 11, 1011–1036 (1999).

15. Rocco, B. R., Sweet, R. A., Lewis, D. A. & Fish, K. N. GABA-Synthesizing Enzymes in Calbindin and Calretinin Neurons in Monkey Prefrontal Cortex. Cereb. Cortex 26, 2191–2204 (2016).

16. Billwiller, F. et al. GABA–glutamate supramammillary neurons control theta and gamma oscillations in the dentate gyrus during paradoxical (REM) sleep. Brain Struct. Funct. 1, 3 (2020).

17. Gentleman, R. C. et al. Bioconductor: open software development for computational biology and bioinformatics. Genome Biol 5, R80 (2004).

18. Collaboration, O. S. Estimating the reproducibility of psychological science. Science (80-.). 349, aac4716–aac4716 (2015).

19. Package ‘nlme’. https://bugs.r-project.org (2020).

